# Architectural trade-offs between environmental stability and genomic redundancy reveal divergent pneumoviral entry strategies

**DOI:** 10.64898/2026.08.18.745394

**Authors:** Jianbing Ma, Hui Zhai, Wenxiang Yu, Jinyue Wang, Jie Deng, Lei Wang, Rui Feng, Liang Xue, Enmei Liu, Xiangxi Wang

## Abstract

Human respiratory syncytial virus (RSV) and human metapneumovirus (hMPV) exhibit distinct seasonal epidemiology, with RSV circulating in early autumn and hMPV peaking in midwinter, yet the structural basis for this niche partitioning remains undefined. Here, we integrate *in situ* cryo-electron tomography and functional virology to decode the architectural logic governing their entry dynamics. RSV employs a matrix (M)-regulated prefusion F (pre-F) organization, partitioning trimers into stabilizing hexagonal superlattices and fusion-competent pools to maintain superior thermotolerance. By contrast, hMPV compensates for its intrinsically unstable, monomeric pre-F with extreme ribonucleoprotein polyploidy, packaging ∼4-fold more genome equivalents to ensure productive infection. Fusion events localize exclusively to M-depleted, non-arrayed membrane regions, establishing a spatial checkpoint for activation. These findings reveal a conserved trade-off between environmental resilience and genomic redundancy that dictates divergent pneumoviral entry strategies, explaining their distinct seasonal ecological niches.

## INTRODUCTION

Human respiratory syncytial virus (RSV) and human metapneumovirus (hMPV), the two leading causative agents of acute lower respiratory tract infections in children worldwide, impose a substantial global health burden, accounting for millions of hospitalizations and tens of thousands of deaths annually ^1–3^. While the recent licensure of RSV vaccines and prophylactic monoclonal antibodies marks a milestone in clinical management ^1,4^, no approved vaccines or antivirals yet exist for hMPV, leaving a critical unmet medical need. Epidemiological surveillance across decades and geographic regions has consistently revealed a striking temporal segregation between the two pneumoviruses: RSV epidemics reliably initiate in late autumn, whereas hMPV activity typically peaks in mid-winter to early spring ^5,6^. This robust phenological lead of RSV suggests intrinsic differences in environmental persistence, transmission fitness, and invasion strategies between the two closely related pathogens. Yet, despite their high sequence and structural homology, the molecular and architectural bases for these divergent clinical patterns remain poorly understood, representing a key gap in our understanding of pneumoviral pathogenesis.

Both RSV and hMPV are enveloped, non-segmented negative-strand RNA viruses that assemble into pleomorphic virions, with filamentous particles representing the dominant morphotype in infected airway cultures ^7–9^. The matrix (M) protein forms a continuous lattice beneath the viral envelope, serving as a geometric scaffold that coordinates the positioning of the fusion glycoprotein (F) and the packaging of helical ribonucleoprotein (RNP) complexes ^8,10^. The prefusion conformation of F (pre-F) is the primary driver of membrane fusion and the principal target of potent neutralizing antibodies and licensed RSV vaccines ^1,8,11–15^. While recent studies have shed light on static snapshots of individual viral components ^10,14,16–19^, the dynamic process of pneumoviral entry remains largely enigmatic. A central unanswered question is how the transition from rigid, filamentous particles, which favor persistent cell-to-cell spread, to fusion-competent, spherical morphotypes is regulated, and what roles the M lattice and F oligomeric states play in this remodeling. Adding to this complexity, an *in vitro* structural study of a prefusion-stabilized hMPV F ectodomain revealed a unique propensity for monomeric prefusion-like conformations ^16^, a feature absent in RSV, but whether this oligomeric plasticity exists in native virions and how it impacts fusion dynamics remains unknown. These gaps hinder our ability to link viral architecture to functional outcomes, including the differential environmental stability that underpins the distinct seasonal ecologies of RSV and hMPV.

Addressing these questions requires structural characterization of native virions in a near-physiological state, minimizing artifacts introduced by conventional purification workflows involving centrifugation, concentration, and buffer exchanges that disrupt labile protein-protein interactions and trigger unintended conformational changes ^8,9,20,21^. Cryo-electron tomography (cryo-ET) of freshly budded, unpurified virions has emerged as a transformative tool for visualizing viral architecture *in situ*, preserving native protein assemblies and transient intermediates ^8,22^. Prior applications of this approach have resolved the helical architecture of RSV RNPs, the dimer-of-trimers organization of RSV pre-F, and the M lattice of related paramyxoviruses ^8,20,23,24^, but no study has yet performed a systematic, side-by-side comparison of RSV and hMPV to link structural divergence to phenotypic differences. Here, we integrate high-throughput cryo-ET, subtomogram averaging, and functional virology to define the architectural logic of RSV and hMPV virions, and to dissect how environmental cues reshape their envelope organization to regulate fusion competency. Our analyses reveal that RSV and hMPV have evolved divergent structural solutions to balance the conflicting demands of environmental resilience and invasive potency: RSV maintains a tightly regulated M-F interface that partitions pre-F trimers into stabilizing superlattices and fusion-ready pools, underpinning its superior thermotolerance and early seasonal emergence, while hMPV compensates for its intrinsically unstable F protein with extreme RNP polyploidy and a low-constraint monomeric F surface layer optimized for rapid fusion triggering. These redefine our understanding of class I fusion protein regulation, explain the structural basis for the distinct epidemiological profiles of RSV and hMPV, and open new avenues for vaccine and antiviral development targeting the dynamic viral envelope.

## RESULTS

### Superior environmental stability underpins the earlier seasonal emergence of RSV relative to hMPV

RSV and hMPV exhibit distinct yet overlapping seasonal epidemiology ^5^. Analysis of multi-year syndromic surveillance data across diverse geographic locales revealed a consistent temporal segregation ^5,6^: RSV epidemics reliably initiated in the late autumn, whereas hMPV activity typically peaked in mid-winter to early spring (Figure S1A). Quantification of epidemic onset timing demonstrated that RSV circulation preceded hMPV in 90.9% of documented seasons (Figure S1B). This robust phenological lead suggests intrinsic differences in viral environmental persistence and transmission fitness under ambient conditions, potentially reflecting divergent structural integrity at the virion level.

To test the hypothesis that RSV possesses greater physicochemical resilience, we performed comparative thermal inactivation kinetics. Viruses were propagated in both Vero-E6 and HEp-2 cells, the latter representing the historical standard for RSV quantification ^12,25^, with both systems yielding comparable cytopathic effects and thermal stability profiles, ensuring assay robustness (Figure S1C). We subjected virus-containing supernatants to acute heat stress (10 min) across a gradient of physiologically relevant temperatures (See Methods). While both viruses exhibited temperature-dependent decay in infectivity, RSV demonstrated markedly superior thermotolerance. At 55 °C, RSV titers declined by approximately 22-fold relative to the 37 °C control, whereas hMPV infectivity plummeted by over 873-fold (Figure 1A). This disparity was exacerbated at 57 °C, where RSV retained detectable infectious particles (192-fold reduction), while hMPV infectivity dropped below the limit of detection (>30,000-fold reduction). These data establish RSV as one of the most thermotolerant enveloped respiratory viruses characterized to date.

**Figure 1.**
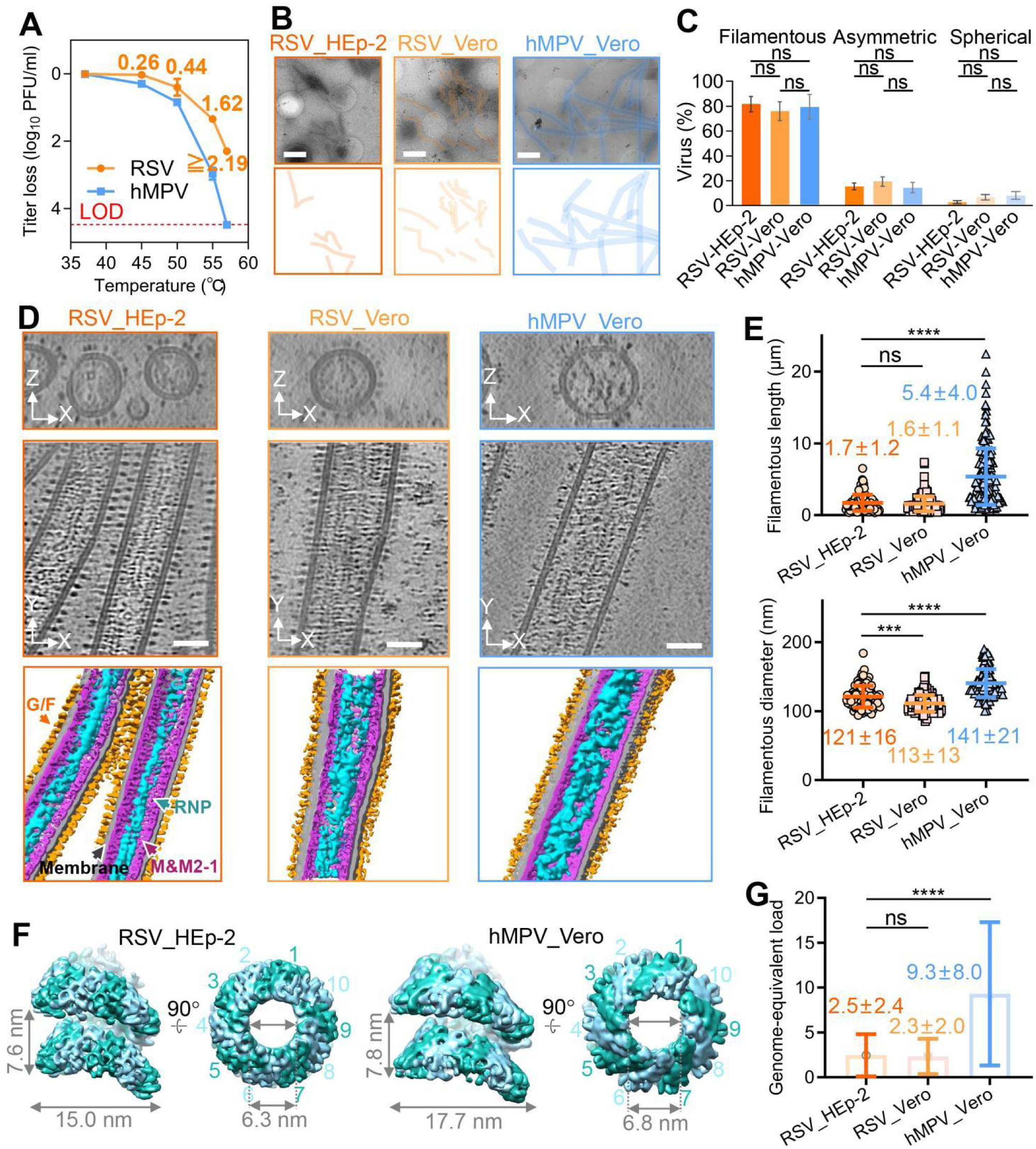
RSV and hMPV assemble distinct envelope-RNP architectures in freshly budded virions. (A) Infectious-titer loss of RSV and hMPV after 10 min of incubation at the indicated temperatures. Incubation at 37°C was used as the low-temperature reference condition. Data are mean ± SD. (B) Representative cryo-EM micrographs of freshly released RSV particles produced in HEp-2 or Vero cells and hMPV particles produced in Vero cells. Filamentous virions are highlighted. Scale bars, 1 μm. (C) Quantification of virion morphology. Particles were classified as filamentous, asymmetric, or spherical. Statistical significance was determined by two tailed unpaired t-test analysis. (D) Representative central slices from cryo-electron tomograms of filamentous virions and corresponding segmentations. The viral membrane is shown in gray, surface glycoprotein (G/F) densities are shown in orange, the matrix (M) layer and putative M2-1-associated internal densities are shown in purple, and ribonucleoprotein (RNP) complexes are shown in blue. Scale bars, 50 nm. (E) Quantification of particle dimensions for RSV and hMPV produced in the indicated cell lines. Data are mean ± SD. Statistical significance was determined by two-tailed unpaired t test; *** p < 0.001, **** p < 0.0001. (F) Subtomogram averages of native helical RNP complexes from RSV and hMPV. Alternating RNP subunits are colored light and dark blue. (G) Estimated RNP-associated genome-equivalent content per filamentous virion. Genome-equivalent content was calculated from RNP helical parameters, RNP contour length per length of virion, and viral length. Nucleotide density was estimated using seven nucleotides per N protein, based on previous RNP structures of RSV and hMPV (PDB: 8OP1 and 8PDN). Data are mean ± SD. Statistical significance was determined by two-tailed unpaired t test; **** p < 0.0001. See also Figure S1 and S2.

### Distinct envelope-RNP architectures underlie differential physicochemical resilience of RSV and hMPV

To elucidate the structural basis for the divergent environmental stabilities of RSV and hMPV, we employed cryo-ET to visualize freshly budded virions in a near-native state (Figures 1B-D and S2A). While both viruses predominantly adopted filamentous morphologies, consistent with previous observations of RSV *in situ* ^8,9,23^, quantitative morphometry revealed profound architectural distinctions. RSV filaments exhibited relatively constrained dimensions, averaging 1.7 ± 1.2 μm (HEp-2) and 1.6 ± 1.1 μm (Vero), with diameters of 121 ± 16 nm and 111 ± 13 nm, respectively. In contrast, hMPV produced markedly elongated and wider filaments, averaging 5.4 ± 4.0 μm in length (approximately 3.3-fold longer than RSV) and 141 ± 21 nm in diameter, with extreme instances exceeding 20 μm (Figures 1B-E). This significant increase in the virion extent and envelope area in hMPV may render its envelope more susceptible to environmental insults, potentially explaining its lower thermotolerance relative to RSV.

Further subtomogram averaging refined the organization of the internal RNP complexes. RSV RNPs exhibited a canonical helical symmetry with 10 nucleoproteins (N) per turn and a pitch of 7.6 nm (Figures 1F, S2B, S2C). Notably, this native architecture diverges sharply from the non-canonical “super-helix” (16 N/protomers per asymmetric unit spanning ∼1.5 turns) recently reported for recombinant RSV N-RNA assemblies analyzed *in vitro*, where inter-turn stabilization is mediated by the C-terminal arm (CTD-arm) of N ^10^. These results suggest that within the confined environment of the virion, constraints imposed by the overlying M lattice and other virion components may disfavor the adoption of this non-canonical super-helical structure (Figure 1D). Instead, the RNP adopts a canonical helical architecture within the native virion. hMPV RNPs displayed a conserved helical organization (10 N/protomer, 7.8 nm pitch), yet possessed slightly larger inner and outer diameters (6.8 nm and 17.7 nm) compared to RSV (6.3 nm and 15.0 nm) (Figure 1F). This structural conservation highlights a conserved nucleocapsid packaging strategy within the *Pneumovirinae*, while the dimensional differences likely reflect species-specific variations in N-protomer geometry and lateral N–N interfaces.

The most striking divergence lay in the estimated genome-equivalent content per particle. Leveraging the determined helical parameters and measured filament lengths, we calculated the RNP content. While RSV virions contained an average of 2.5 ± 2.4 (HEp-2) and 2.3 ± 2.0 (Vero) genome equivalents, aligning with prior reports of RSV polyploidy ^20,25,26^, hMPV filaments harbored a remarkably high genome equivalents per particle (Figures 1G, S2D). This ∼3.9-fold increase in genomic payload suggests that hMPV relies on a high-copy-number strategy for genome delivery. Functionally, this polyploidy may buffer against the degradation of genomic RNA during its transit through the harsh extracellular environment or provide a competitive advantage during the early stages of infection by saturating the host’s innate immune sensors with excess viral RNA or proteins ^2,25,27,28^. However, the energetic cost of packaging such lengthy RNPs, coupled with the fragility of the expansive envelope, may underpin the observed trade-off between hMPV’s high intracellular genomic capacity and its comparatively limited environmental stability relative to RSV.

### M-dependent spatial segregation of RSV pre-F into dimers-of-trimers and hexagonally packed superlattices

The pre-F of the RSV fusion glycoprotein is the principal target of neutralizing antibodies and the cornerstone of licensed vaccines ^14,29,30^. While pre-F is known to organize as dimers-of-trimers on filamentous virions ^9,20,23^, the regulatory mechanisms governing its higher-order assembly remain obscure. Here, using high-throughput cryo-ET and subtomogram averaging of freshly budded virions, we resolve two biochemically distinct pre-F organizational states dictated by the underlying M protein scaffold (Figure 2).

**Figure 2.**
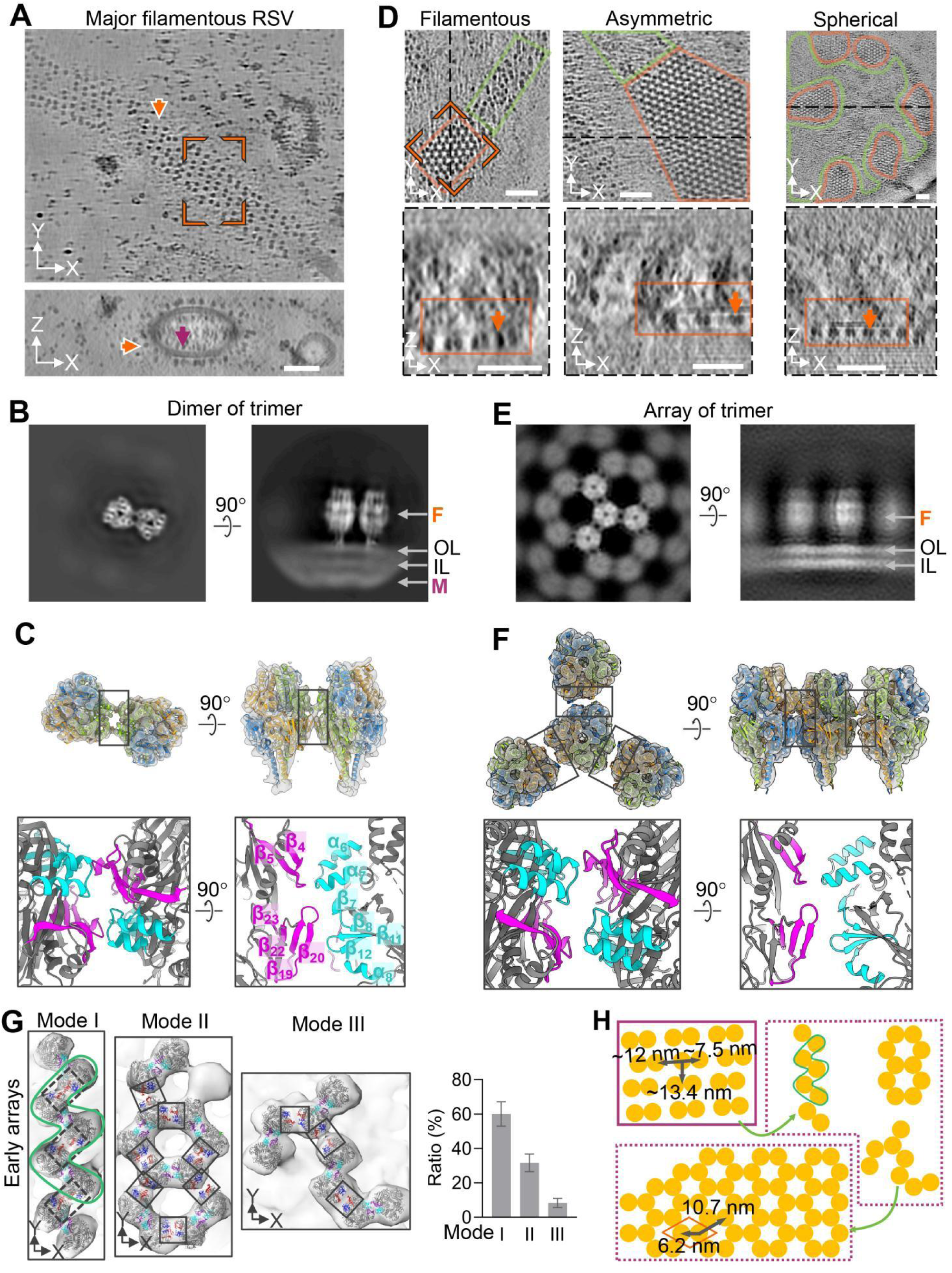
RSV pre-F forms M-associated dimers of trimers and M-depleted hexagonal arrays in situ. (A) Representative cryo-ET surface slices of filamentous RSV particles. Surface pre-F densities are shown in orange, and the M layer beneath the viral membrane is shown in purple. Paired pre-F densities corresponding to dimers of trimers are indicated. Scale bars, 50 nm. (B) Representative slices through the subtomogram average of RSV pre-F dimers of trimers reconstructed with the viral membrane included. The outer leaflet (OL), inner leaflet (IL), and underlying M density are indicated. (C) Subtomogram average of RSV pre-F dimers of trimers reconstructed without membrane signal. The density fit well with the RSV pre-F trimer model (PDB: 4MMS). The inter-trimer interface is boxed, with enlarged views shown below; interface-proximal elements are highlighted in purple and cyan. (D) Representative cryo-ET surface slices showing hexagonal pre-F arrays on filamentous, asymmetric, and spherical RSV particles. Ordered arrays are boxed in orange, adjacent regions with reduced or less ordered pre-F density are boxed in green. Scale bars, 50 nm. (E) Representative slices through the subtomogram average of the hexagonal pre-F array reconstructed with the viral membrane included. (F) Focused subtomogram average of the central pre-F trimer within the hexagonal array, reconstructed without membrane signal. The density fit well with the RSV pre-F trimer model (PDB: 4MMS). Inter-trimer interfaces are boxed, with enlarged views shown below. (G) Representative early array-assembly intermediates of RSV pre-F. Pre-F dimers of trimers were modeled using PDB: 4MMS. Elements near the dimer interface are highlighted in purple and cyan, whereas elements near interfaces between adjacent dimers of trimers are highlighted in red and blue. Interfaces are indicated by solid or dashed outlines. One M-shaped unit within the larger assembly is outlined in green. (H) Model for RSV pre-F higher-order assembly, linking M-associated dimers of trimers, M-depleted hexagonal arrays, and progressive engagement of lateral inter-trimer interfaces. See also Figure S3.

In the canonical state, pre-F predominantly organized as dimers-of-trimers (Figure 2A). Subtomogram averaging of 40,544 particles yielded a 9.4 Å reconstruction (C1 symmetry), which sharpened to 6.9 Å upon application of C2 symmetry (Figures 2B, 2C, S3A-S3C). The unambiguous fit of the atomic model (PDB: 4MMS) confirmed the native prefusion state ^14^. Crucially, a continuous M protein layer was resolved directly beneath these dimeric assemblies (Figures 2B, S3A), corroborating recent findings that the M lattice serves as a geometric template for F positioning ^8^. The dimer interface was stabilized by extensive contacts involving the α_6_-α_7_, β_4_-β_5_ and β_19_-β_20_ regions, effectively occupying one of the three geometrically permissible lateral interfaces surrounding each trimer (Figures 2C).

Strikingly, we identified a previously unresolved, highly ordered hexagonal lattice of pre-F on subsets of virions (Figures 2D-F, S3D). This “superlattice” exhibited sharp Fourier reflections, indicative of long-range two-dimensional crystalline order. Subtomogram averaging of 15,280 particles yielded an 11.0 Å reconstruction, refined to 7.4 Å with C3 symmetry focusing on the central trimer (Figures 2F, S3D-S3H). The atomic model fitting confirmed that this lattice consists exclusively of F trimers in the prefusion conformation. In contrast to the canonical state, the density corresponding to the M layer was conspicuously absent beneath these hexagonal arrays (Figures 2B, 2E). This spatial segregation indicates that the formation of extensive pre-F superlattices is facilitated by, or perhaps induces, local M protein dissociation.

The transition from dimers-of-trimers to hexagonal arrays represents a shift from partial to full interface occupancy. In the dimeric state, only one lateral interface per trimer is engaged; in the hexagonal lattice, all three interfaces are engaged, promoting cooperative assembly (Figures 2C, 2F). We captured transient assembly intermediates that elucidate this maturation process (Figures 2G, S3I). Mode I intermediates (60 ± 7% of events) displayed an “M-shaped” trimer configuration, suggesting a nucleation event preceding lattice expansion. Modes II and III exhibited interface geometries more congruent with the mature lattice (Figures 2G). Quantitative analysis revealed that the hexagonal lattice (20,026 trimers/μm^2^) is significantly denser than the dimer-of-trimers configuration (7,654 trimers/μm^2^) (Figures 2H and S3J, S3K). Membrane regions immediately adjacent to mature arrays were frequently devoid of pre-F density (Figure 2D, green boxes), supporting a model of recruitment-driven coalescence, wherein trimers migrate laterally to join the expanding lattice, leaving F-depleted “halos”.

### Quaternary instability of hMPV pre-F underpins its disordered surface architecture and thermal susceptibility

In contrast to the highly ordered pre-F superstructures observed on RSV, hMPV filaments exhibited a sparse and heterogeneous glycoprotein layer (Figures 1D and 3A). While nearest-neighbor distance analysis revealed a lateral spacing (8.6±2.0 nm) comparable to that of RSV pre-F assemblies (Figures S3J and S4A), the apparent size and density of individual hMPV glycoprotein spikes were markedly reduced (Figures 3A). To define the oligomeric state of hMPV F *in situ*, we performed targeted subtomogram averaging guided by an RSV pre-F template. This analysis resolved two distinct populations: a minor fraction (14%) compatible with a trimeric architecture at 32.1 Å, and a dominant fraction (86%) whose mass and dimensions corresponded to monomeric F at 32.4 Å (Figures 3B, S4B, S4C). These findings demonstrate that, unlike RSV, native hMPV virions predominantly display F in a protomeric state rather than a stable prefusion trimer.

**Figure 3.**
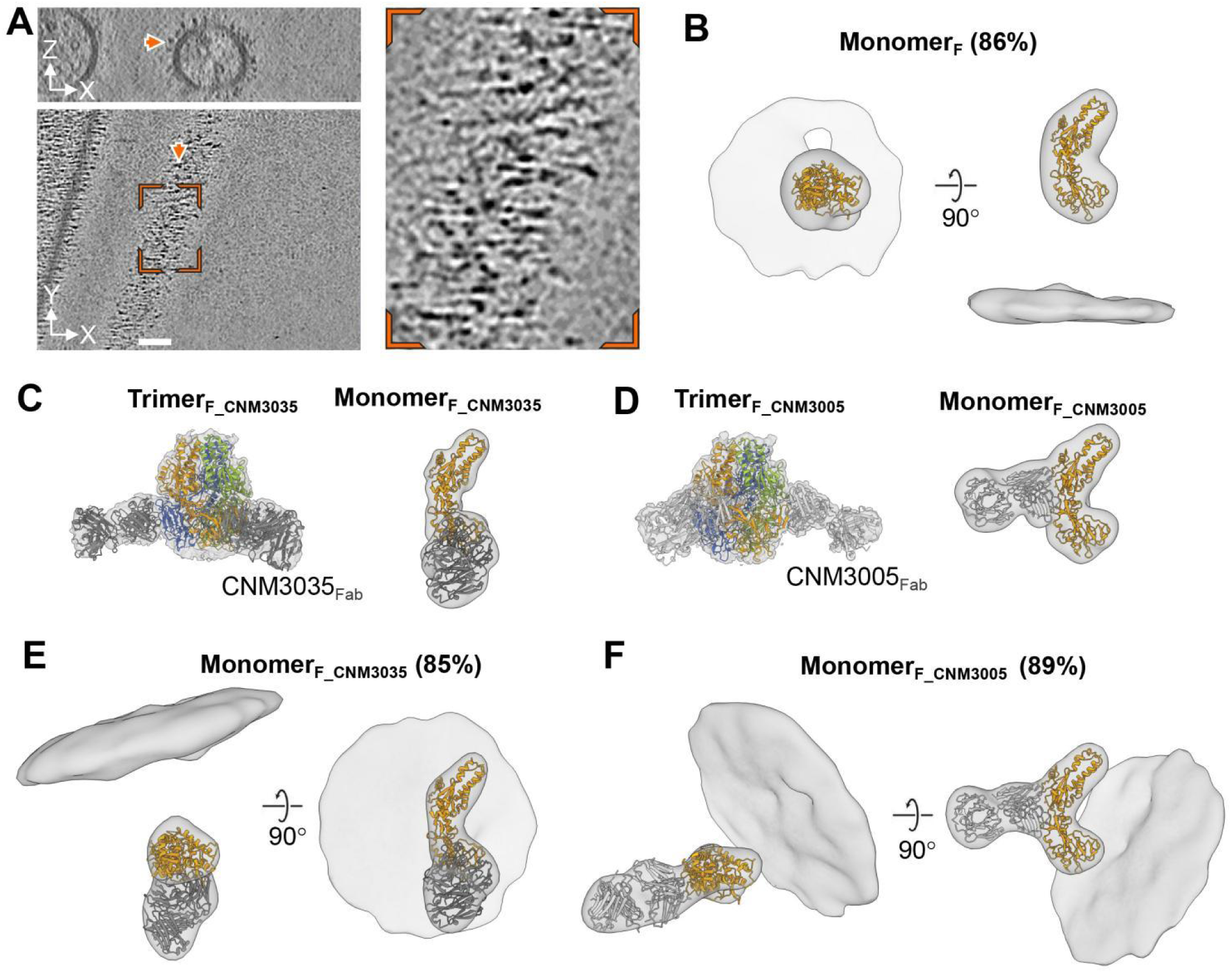
hMPV pre-F is predominantly monomeric on native virions. (A) Representative cryo-electron tomographic surface slice of a filamentous hMPV particle. Surface glycoprotein densities are highlighted in orange, and an enlarged view of representative surface densities is shown. Scale bars, 50 nm. (B) Subtomogram average of hMPV surface F densities on native virions. The density is compatible with a monomeric hMPV pre-F model, shown in orange. The fraction of particles assigned to the monomeric pre-F class is indicated. (C and D) Surface representations of single-particle cryo-EM reconstructions of purified hMPV pre-F trimers in complex with CNM3035 (C) or CNM3005 (D) Fab fragments. Cryo-EM maps are shown in light gray, hMPV pre-F protomers are colored orange, green, and blue, and Fab models are shown in dark gray (C) or light gray (D). (E and F) Subtomogram averages of hMPV surface F densities on intact virions after incubation with CNM3035 (E) or CNM3005 (F) Fab fragments. The densities are compatible with monomeric hMPV pre-F bound to Fab, with F shown in orange and Fab density shown in dark gray (E) or light gray (F). The fractions of particles assigned to the Fab-bound monomeric pre-F classes are indicated. See also Figure S4.

To validate the molecular identity of these densities, we generated Fab fragments from two hMPV-specific neutralizing mAbs (CNM3035 and CNM3005) isolated from convalescents ^31^. Structural characterization by single-particle cryo-EM confirmed that both Fabs bound monomeric and trimeric hMPV F *in vitro* (3.6-3.7 Å for trimers; 10.3-17.6 Å for monomers; Figures 3C, 3D, S4D, S4E). Crucially, epitope mapping revealed that binding was confined to individual F protomers, obviating the need for quaternary interfaces. This property allowed these two Fabs to serve as stoichiometric probes for F irrespective of its oligomeric state. Applying these probes to intact virions, we found that hMPV F remained overwhelmingly monomeric (85-89% of particles; at 26.6-29.2 Å; Figures 3E, 3F, S4B). Minor trimeric populations were detected (at 27.7-31.1 Å; Figures S4F, S4G), but even under saturating conditions, steric constraints within the crowded viral envelope precluded the binding of a third Fab per trimer, as the inter-spike distance (8.6 nm) was insufficient to accommodate the ∼14 nm span of a Fab-decorated trimer. This quaternary instability of hMPV F structurally explains the weak, disordered surface density observed in cryo-ET, as the absence of stable inter-trimer interactions prevents the formation of cohesive lattice structures seen in RSV (Figure 2). Consequently, the propensity of F to dissociate into monomers likely lowers the kinetic barrier for thermal denaturation, underpinning the heightened thermosensitivity of hMPV virions (Figure 1). Furthermore, the inherent structural fragility presumably necessitates a compensatory strategy of packaging higher genome equivalents to ensure productive infection, illustrating a critical trade-off between envelope stability and genomic redundancy in pneumoviruses.

### Matrix disassembly triggers cooperative pre-F lattice formation and enhances thermotolerance

Prior studies reveal that environmental stressors including heat and purification-associated mechanical shear induce coordinated structural rearrangements in RSV virions, including prefusion-to-postfusion conversion of F-trimer, disruption of the ordered M layer, and remodeling of filamentous particles into spherical morphologies ^20,21^. Whether these changes represent mechanistically coupled steps or parallel consequences of generalized virion destabilization has remained unresolved. Our cryo-ET analyses have shown that the higher-order organization of RSV F does not mirror the periodicity of the M lattice, implying that coupling between the two layers is mediated by local, rather than long-range, interactions (Figure 2). Building on this, we used IsoNet missing-wedge-corrected tomography to resolve the structural linkage between F and M ^32^. We found that the C3-symmetric ectodomain of pre-F is not propagated to its membrane-proximal region: 81 ± 8% of pre-F trimers exhibited splayed, monomer-like transmembrane-cytoplasmic tail (TM-CT) densities that extended individually to tether to the underlying M layer, while only 19 ± 4% displayed compact, trimer-like TM-CT bundles associated with the matrix (Figures 4A, S5A, S5B). Both modes of F anchoring indicate that the intact M lattice imposes steric constraints on the viral envelope that presumably restrict the mobility of pre-F, maintaining the structural rigidity of the virion membrane under baseline conditions.

**Figure 4.**
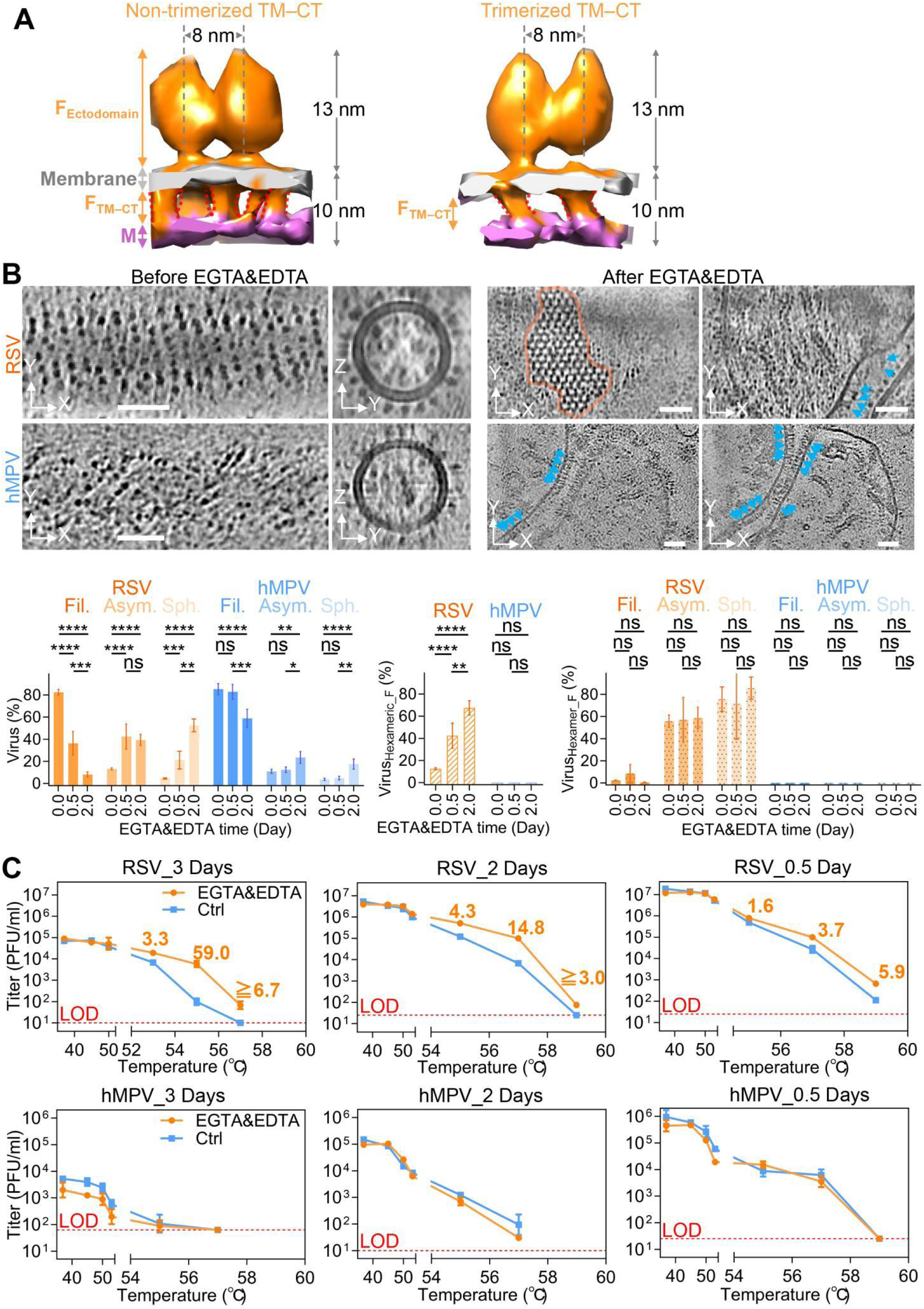
Matrix remodeling promotes RSV pre-F array formation and virion thermotolerance. (A) Surface representations of typical subtomogram maps of RSV pre-F dimers of trimers with membrane and M densities retained. The TM–CT region shows predominantly non-trimeric organization (left) and a minor trimeric organization (right). (B) Quantification of virion morphology and surface-assembly states in RSV and hMPV before and after EGTA & EDTA treatment. Ordered arrays are in orange. The post-like F are indicated by cyan arrows. Data are mean ± SD from two independent experiments. Statistical significance was determined by two-tailed unpaired t test; * p < 0.05, ** p < 0.01, *** p < 0.001, **** p < 0.0001. Scale bars, 50 nm. (C) Infectious titers after 10 min of incubation at the indicated temperatures. Before heat treatment, virions were incubated with EGTAand EDTAfor the indicated times and then supplemented with CaCl_2_ and MgCl_2_ before infectivity measurement. Data are mean ± SD from two independent experiments. LOD, limit of detection. See also Figure S5.

Given that M dimerization and higher-order assembly are strictly dependent on binding of divalent cations ^17,18^, we reasoned that depletion of cations would selectively disrupt the M scaffold without introducing confounding thermal stress, providing a route to dissect the causal relationship between M organization and F dynamics. We treated freshly budded virions with 5 mM EGTA & EDTA, and analyzed structural changes by cryo-ET (Figure 4B). Chelation induced a dramatic, coordinated remodeling of RSV architecture over 0.5–2 days: filamentous particles decreased from 83 ± 3% to 8 ± 3% of the total population, while spherical particles surged from 4 ± 1% to 53 ± 6%, and asymmetric particles rose from 13 ± 2% to 39 ± 5% (Figure 4B). Concurrent with this morphological shift, the proportion of particles containing ordered hexagonal pre-F arrays increased nearly sixfold, from a baseline of 13 ± 1% to 68 ± 7% (Figures 4B, S5C-S5E). Critically, these arrays were almost exclusively localized to regions of the viral envelope that lacked underlying M density (Figures S5E), confirming that disassembly of the M lattice relieves the steric constraints that normally inhibit lateral F mobility, permitting the cooperative diffusion and repacking of F trimers into long-range ordered hexagonal lattices. This remodeling is RSV-specific: under identical chelation conditions, hMPV underwent partial morphological transitions with filamentous particles decreasing from 85 ± 6% to 59 ± 9%, but failed to assemble hexagonal pre-F arrays (Figure 4B). This deficit aligns with our earlier observation that hMPV F exists predominantly as unstable monomers on native virions, lacking the stable trimeric building blocks required for cooperative lattice formation.

The spatial segregation of F populations within remodeled RSV virions revealed a link between supramolecular organization and conformational stability. F trimers incorporated into hexagonal arrays retained a compact, prefusion-compatible architecture, while regions of the envelope excluded from these arrays frequently contained elongated, postfusion-like densities (Figures 2D, 4B, S3E). This dichotomy reflects two countervailing consequences of M disassembly. Trimers embedded in hexagonal lattices are fully saturated by neighboring ones, forming a multivalent network that kinetically traps and stabilizes the metastable prefusion conformation, raising the energy barrier for spontaneous structural rearrangement. By contrast, F trimers excluded from these lattices lose M-mediated anchoring, gaining higher conformational flexibility and a drastically lowered energy threshold for pre-to-post fusion transition. Notably, this structural arrangement does not preclude infectivity: rather, it partitions F trimers into two functional pools, one stabilized for environmental persistence and one primed for efficient fusion initiation upon host cell encounter.

To test whether this structural partitioning translates to enhanced fitness, we measured the thermotolerance of array-enriched RSV populations, replenishing Ca^2+^/Mg^2+^ prior to heat challenge to control for acute cation depletion effects. At temperatures up to 50 °C, array-enriched and untreated virions retained comparable infectivity, indicating that matrix remodeling and array formation do not impair entry competence under permissive conditions. Above 50 °C, however, array-enriched RSV exhibited dramatically improved survival: infectivity retention was enhanced 1.6-fold after 0.5 days of chelation, 4.3-fold after 2 days, and 59.0-fold after 3 days of pretreatment (Figure 4C). No comparable protection was observed for hMPV, consistent with its inability to form stabilizing pre-F arrays. These findings reveal that M-lattice disassembly initiates a hierarchical remodeling cascade in RSV: loss of M-mediated tethering permits F lateral diffusion and hexagonal lattice formation, stabilizing the prefusion conformation, while non-arrayed regions retain a pool of fusion-competent F trimers. This dual-functional organization provides a mechanistic explanation for RSV’s superior thermotolerance relative to hMPV, and resolves the long-standing question of how stress-induced structural changes in RSV are causally coupled through the envelope-matrix interface.

### Hexagonal pre-F arrays coincide with elevated fusion frequency outside ordered lattices

To determine whether M-layer remodeling and the concomitant formation of hexagonal pre-F arrays alter the propensity for fusion activation, we quantified membrane-apposition events and F-associated conformational intermediates across RSV tomograms (Figure 5A). Consistent with the morphological shifts induced by chelation, fusion-associated features were rarely observed on intact filamentous particles (0.26±0.15%). In contrast, these events were significantly enriched on remodeled virions, occurring at 2.46 ± 0.99% of asymmetric particles and 4.80 ± 1.82% of spherical particles, an approximately 9.5-fold and 18.5-fold increase, respectively (Figure 5B). This enrichment reveals that virion remodeling promotes a biophysical environment conducive to membrane-proximal F interactions. We identified 27 distinct fusion-associated sites involving viral membranes and adjacent vesicles or cellular debris (Figure 5C, 5D). These sites yielded two characteristic structural classes at limited resolution. The first class featured extended densities (∼20–24 nm) consistent with the dimensions of a prehairpin intermediate bridging adjacent bilayers (Figure 5C). The second class comprised pore-like membrane continuities associated with shorter densities (∼17 nm) compatible with postfusion F (Figure 5D). While the resolution precludes atomic modeling, the dimensions and spatial organization of these densities are consistent with canonical RSV F refolding intermediates ^33,34^.

**Figure 5.**
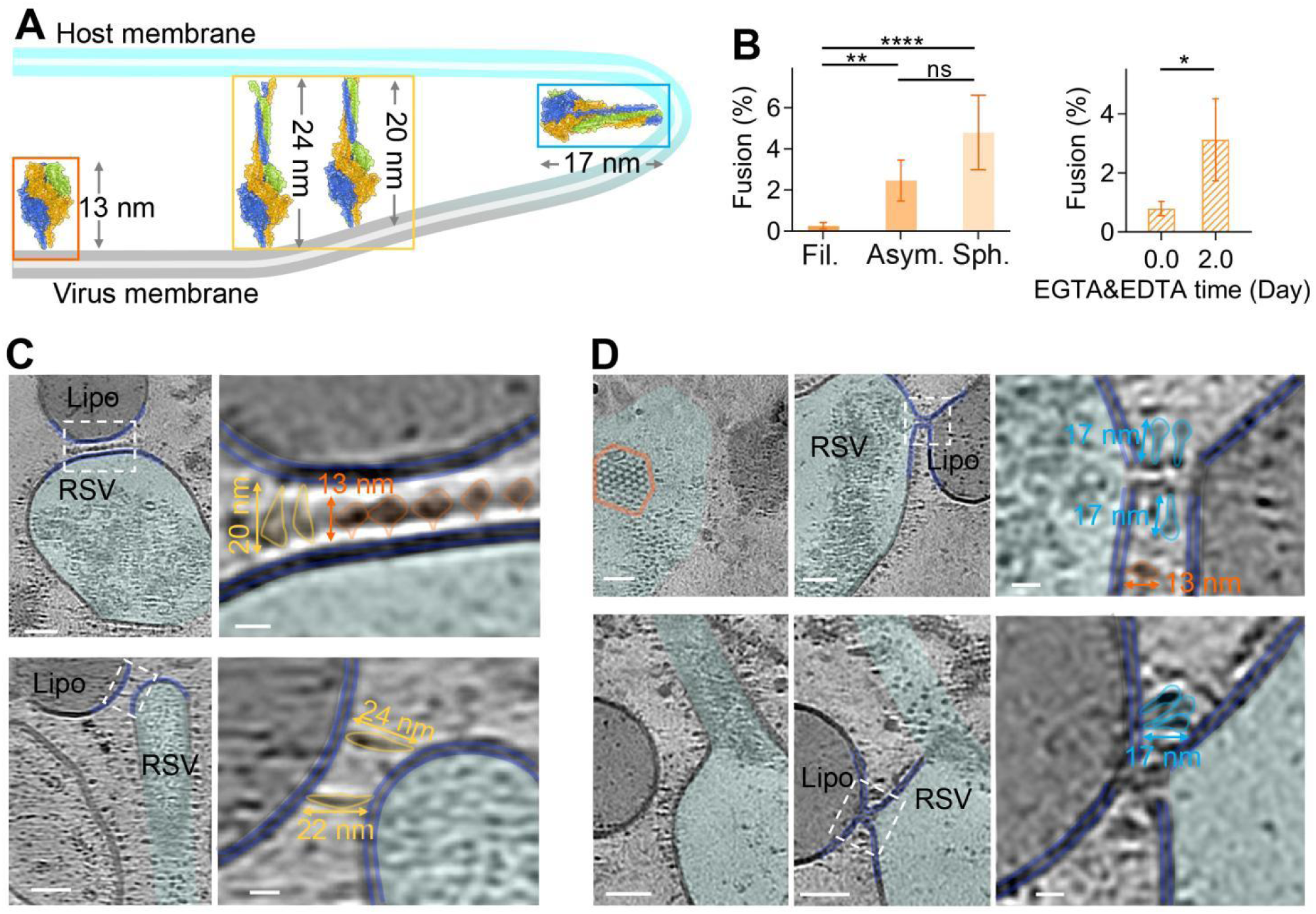
Fusion-associated intermediates are captured on RSV particles. (A) Structural model of RSV F conformational transitions during membrane fusion. Prefusion F, extended prehairpin-like intermediate, and postfusion F are shown with approximate molecular lengths indicated. The model was generated using RSV prefusion F and postfusion F structures (PDB: 4MMS and 6APB). (B) Quantification of fusion cases in RSV. Data are mean ± SD. Statistical significance was determined by Fisher’s exact test; * p < 0.05, ** p < 0.01, **** p < 0.0001. (C) Representative tomographic slice showing an RSV particle with a membrane-bridging fusion-associated intermediate. The boxed region is enlarged on the right to highlight extended F-like densities connecting the viral and apposed membranes. Viral membranes are outlined in blue, and matrix density beneath the viral membrane is highlighted in purple. Lipo, cell-derived liposome. Scale bars, 50 nm. (D) Representative tomographic slices showing RSV particles with fusion-pore-like membrane-continuity events. Boxed regions are enlarged on the right to highlight associated F-like densities compatible with postfusion F. Ordered hexagonally packed pre-F arrays are boxed in orange where present. Scale bars, 50 nm.

Critically, none of these fusion events localized to the interior of intact hexagonal pre-F arrays. All but one occurred in membrane regions lacking underlying M density and spatially separated from ordered lattices; the single exception resided at an incomplete array margin (Figure 5C). This spatial segregation indicates that while M-disassembly and particle rounding coincide with a higher frequency of fusion-competent F conformations, these events are restricted to regions where F trimers are not immobilized within the multivalent lattice. Thus, the formation of hexagonal arrays coincides with an increased probability of observing fusion events elsewhere on the virion envelope, consistent with the model that lateral F–F contacts within the lattice restrict access to the extended intermediates required for membrane fusion. In contrast, hMPV virions yielded only one unambiguous fusion-associated event (Figure S6), reflecting the paucity of ordered F trimers available to support canonical fusion intermediates.

## DISCUSSION

Building upon our structural and functional delineation of RSV and hMPV virions, we propose a dynamic model for pneumoviral entry that reconciles viral morphology, M organization, and fusion activation (Figure 6). Upon initial adherence to the host cell surface, the long filamentous morphology of both viruses, characterized by regularly spaced prefusion F trimers/monomers (7.5 ± 1.0 nm in RSV; 8.6 ± 2.0 nm in hMPV), maximizes avidity for cellular receptors ^35–37^, facilitating stable tethering. This morphological advantage aligns with recent *in situ* cryo-ET studies showing that filamentous RSV particles dominate in infected airway cultures, where their elongated form promotes persistent cell-to-cell contacts ^8,9,38^. Subsequent entry kinetics diverge sharply following this adhesion phase (Figure 6). For RSV, we propose that host factors, putatively including furin-mediated further cleavage to release the p27 peptide (a process previously suggested to be more efficient in spherical versus filamentous morphotypes ^26^ or other undefined enzymatic activities, may trigger the disassembly of the M lattice. This dismantling alleviates the steric constraints on the envelope ^20,23^, converting the rigid filament into a more pliable spherical particle (Figure 6). Besides, the morphological transition liberates F trimers from their rigid M-mediated tethers, permitting their lateral diffusion and coalescence into hexagonal pre-F superlattices. Crucially, these processes partition the F population: the majority is sequestered into stable arrays, while a minority remains dispersed in regions of high membrane fluidity (Figure 6). This spatial segregation is key to RSV’s “dual-mission” strategy—maintaining a reservoir of stable trimers for environmental persistence while generating a pool of fusion-competent, unconstrained trimers poised for membrane fusion (Figure 6). In contrast, hMPV employs a complementary, evolutionarily distinct strategy. While hMPV also adheres efficiently, its intrinsic lack of stable F trimers and ordered F-lattice interactions bypasses the need for complex remodeling. Instead, hMPV leverages its extreme polyploidy (packaging nearly four times the genomic equivalents of RSV) and its longer morphology (Figure 6). The genomic redundancy aligns with recent evidence that pneumoviruses can function as “collective infectious units” where multiple RNPs enter a single cell to facilitate the formation of viral factory, enhancing transcription, replication and immune evasion ^2,27,28^. Coupled with its sparse, monomeric F surface density, the lower spatial and conformational constraints on hMPV F may paradoxically facilitate more rapid or stochastic triggering once anchored to a host cell, compensating for its inferior thermotolerance with a brute-force genomic payload and efficient receptor engagement.

**Figure 6.**
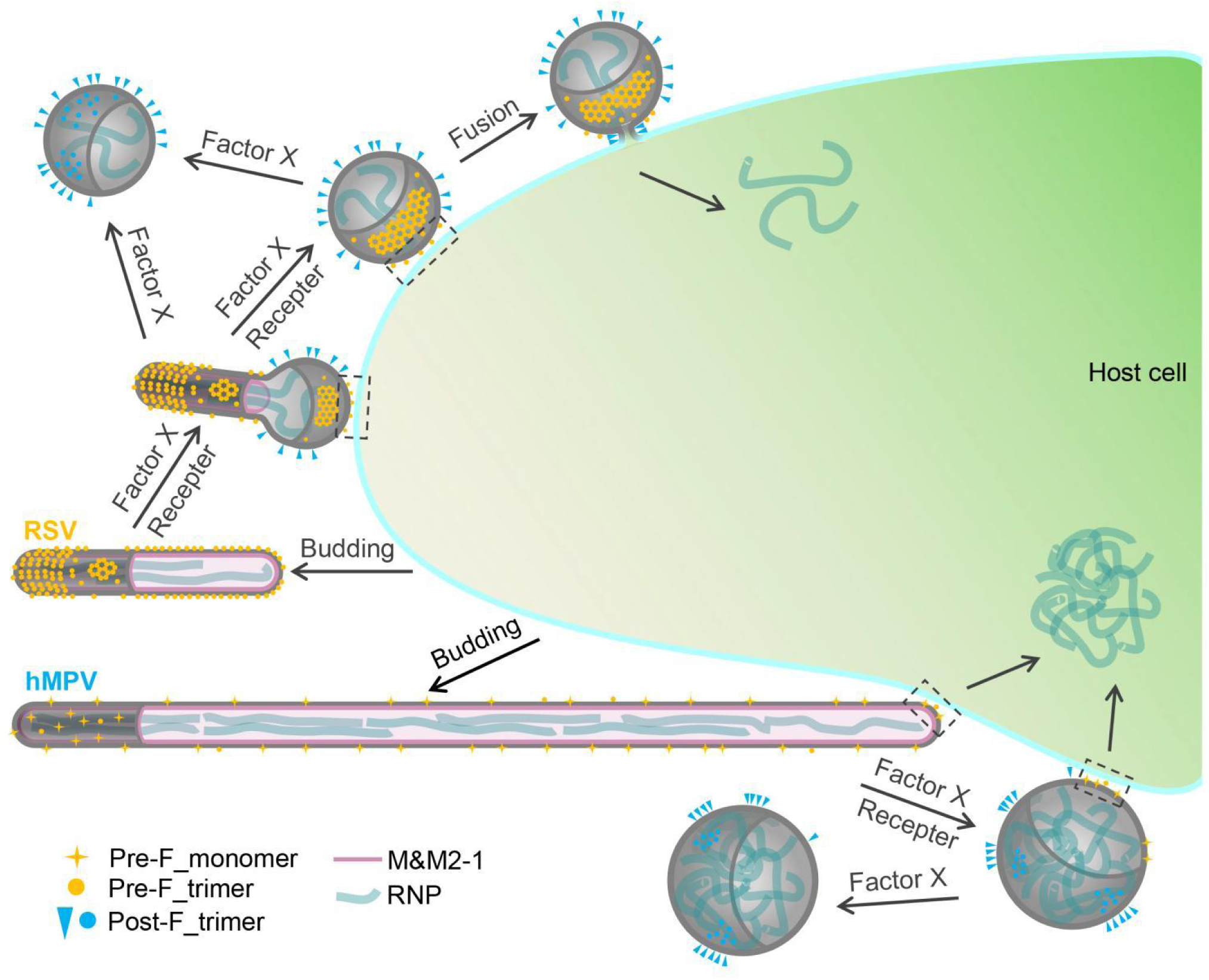
Model for divergent virion-scale remodeling and F-mediated membrane fusion in RSV and hMPV. Newly budded RSV and hMPV particles are predominantly filamentous. Host-associated factors or other perturbations, represented schematically as “Receptor” and “Factor X” in the model above, are proposed to disrupt M organization, thereby relieving M-imposed constraints on pre-F mobility and driving the remodeling of filamentous virions into asymmetric and spherical forms. In RSV, this redistribution allows M-associated dimers of trimers to reorganize into M-depleted hexagonal pre-F arrays, alongside non-array pre-F trimers and post-F-like densities. Pre-F trimers that remain outside the arrays, or that transiently dissociate from them, may be freed from lateral interactions that constrain the prefusion-to-postfusion transition, thereby permitting initiation of the canonical class I fusion pathway, including extension toward the target membrane, fusion-peptide insertion, fold-back into the postfusion trimer, and fusion-pore formation. By contrast, hMPV F is observed predominantly as monomeric, prefusion-like densities, suggesting that membrane fusion may proceed from a noncanonical oligomeric starting state. hMPV F may assemble into prefusion trimers or pass through transient alternative oligomeric intermediates before formation of the postfusion trimer. These distinct F organizations are embedded within broader differences in virion morphology and RNP packaging. The longer filaments and higher genome-equivalent RNP content of hMPV may favor short-range spread across adjacent airway epithelial cells and delivery of a larger RNP load, whereas the ordered trimeric and higher-order pre-F organization of RSV, together with its greater thermotolerance, may contribute to preservation of extracellular infectivity.

The hierarchical remodeling observed in RSV mirrors a conserved evolutionary strategy among class I viral fusion proteins that the transient clustering of fusion-competent trimers to optimize the energetics of membrane fusion ^39,40^. Together with previous studies ^14,41^, our results suggest that the prefusion state of F is intrinsically metastable and requires “stabilization in waiting”, either through M-layer tethering or potentially through transient protein-protein interactions in other viruses ^39,42^, to prevent premature inactivation. The RSV-specific formation of hexagonal arrays represents a sophisticated implementation of this principle, creating a high-density, “release-gated platform” for fusion initiation only after the virion has committed to entry. The “cooperative trigger” model, wherein the energy landscape for F refolding is modulated by lateral interactions and membrane topology, likely extends beyond pneumoviruses. The recent observation of higher-order assemblies of prefusion F trimers in paramyxoviruses, receptor-induced clustering of influenza A HA trimers, and hexagonal lattice packing of influenza C HEF trimers ^39,43–45^ suggest that viruses routinely exploit higher-order glycoprotein organization to toggle between stability and fusogenicity. RSV’s dual-functional organization by using the M lattice to enforce rigidity and the F lattice to enforce stability, thus provides a compelling paradigm for how enveloped viruses manage the precarious balance between environmental resilience and invasive potency.

These structural insights carry immediate implications for antiviral development and vaccine design. The RSV pre-F hexagonal array represents a novel, virally encoded immunogen that presents inter-trimer quaternary epitopes not found on soluble trimers or dimer-of-trimers. While current subunit vaccines (e.g., Arexvy, Abrysvo) use trimeric prefusion F antigens and have demonstrated clinical efficacy ^46,47^, the array’s unique antigenic landscape could be exploited to elicit broadly neutralizing antibodies that recognize conserved patches at the trimer-trimer interface (Figure 2). Conversely, the structural fragility of hMPV F, predominantly as monomers, suggests that immunogens designed to lock F into a stable trimeric conformation may be particularly efficacious ^13,16,48^. This approach would prevent spontaneous dissociation that likely contributes to its thermal lability. Furthermore, the identification of M-layer disassembly as a prerequisite for RSV fusion suggests a novel antiviral vulnerability: small molecules targeting the calcium-binding pockets of M could trap virions in a fusion-incompetent state ^17,18^. Clinically, our findings provide a structural basis for the distinct seasonal ecologies of these viruses. RSV’s ability to maintain stable, fusion-ready arrays allows for earlier autumnal circulation when environmental pressures are milder ^5,6,49^. In contrast, hMPV’s reliance on a high-genomic-copy-number strategy may be more suited to the colder, more stable winter environment ^27,50,51^, where the reduced metabolic activity of host cells and lower environmental microbial competition favor a “stealthy” high-yield infection strategy.

We acknowledge several limitations in the present study. While cryo-ET provides unparalleled snapshots of viral architecture in a near-native state, it captures static moments rather than the dynamic continuum of fusion. The low resolution of fusion intermediates precludes definitive atomic modeling of the prehairpin or postfusion states *in situ*, necessitating future studies using time-resolved cryo-ET or correlative light-electron microscopy to track these transitions in real-time. Additionally, our *in vitro* chelation experiments, while informative, are a proxy for the complex biochemical environment of the host airway surface liquid (ASL). The ASL contains varying concentrations of divalent cations, proteases (like furin), and mucins that collectively influence M-lattice integrity and F processing. Future work should aim to replicate these remodeling events in physiologically relevant models, such as differentiated human airway organoids, to validate the proposed entry model. Finally, while we infer the functional consequences of F oligomeric state on fusion efficiency, direct single-virion fusion assays correlating F organization with pore formation kinetics would strengthen the causal link between structure and infectivity. Recent advances in fluorescence-based fusion assays coupled with super-resolution microscopy may prove instrumental in bridging this gap.

In summary, this study elucidates the structural logic underpinning the differential environmental stability and entry dynamics of RSV and hMPV. We demonstrate that RSV utilizes a tightly regulated M-F interface to partition its surface glycoproteins into stabilizing superlattices and fusion-ready pools, a mechanism that directly contributes to its superior thermotolerance. Conversely, hMPV compensates for its structural fragility with genomic redundancy and a simplified, low-constraint fusion apparatus. By resolving the architecture of the pre-F hexagonal array and its spatial exclusion from fusion sites, we provide a structural framework for understanding how pneumoviruses navigate the conflicting demands of stability and infectivity. These findings not only refine our understanding of viral pathogenesis but also chart new avenues for therapeutic intervention targeting the dynamic envelope of these pervasive human pathogens.

## RESOURCE AVAILABILITY

### Lead contact

Requests for further information and resources should be directed to and will be fulfilled by the lead contact, Xiangxi Wang.

### Materials availability

All requests for resources should be directed to and will be fulfilled by the lead contact, with a completed materials transfer agreement (MTA).

### Data and code availability

The cryo-ET density maps generated in this study have been deposited in the Electron Microscopy Data Bank (EMDB): EMD-82792 (RSV) and EMD-82793 (hMPV) for RNP, EMD-82796 and 82797 for dimer of pre-F trimer from RSV, EMD-82800 and 82803 for hexagonal-lattice of pre-F trimer from RSV, EMD-82804 and 82806 for hexagonal-lattice of pre-F trimer from RSV after EGTA & EDTA treatment, EMD-82808 for pre-F monomer from hMPV, EMD-82807 for pre-F trimer from hMPV, EMD-82810 for pre-F monomer in complex with CNM3035_Fab_ from hMPV, EMD-82809 for pre-F monomer in complex with CNM3035_Fab_ from hMPV, EMD-82813 for pre-F monomer in complex with CNM3005_Fab_ from hMPV, EMD-82811 for pre-F monomer in complex with CNM3005_Fab_ from hMPV. The cryo-EM density maps and structure models generated in this study have been deposited in the EMDB and Protein Data Bank (PDB): EMD-82866 and PDB 44RU for pre-F trimer in complex with CNM3035_Fab_ from hMPV, EMD-82811 for pre-F trimer in complex with CNM3005_Fab_ from hMPV.

This work did not use or generate new code.

Any additional information required to reanalyze the data reported in this paper is available from the lead contact upon request.

## ACKNOWLEDGMENTS

This work was supported by Strategic Priority Research Program (No. XDB1310000 to X.W.); National Science and Technology Major Project (No. 2025ZD01903800 to X.W.); National Natural Science Foundation of China (No. 32325004, T2394482 and 12034006 to X.W.; No. 32371286 to J.M.); Basic Research Program Based on Major Scientific Infrastructures, CAS-JZHKYPT-2021-05, CAS (No. YSBR-010 to X.W.); National Key Research and Development Program (No. 2023YFC2306003 to X.W.; No. 2022YFC2704900 to E.L.); Ministry of Science and Technology of China (No. CPL-1233 to X.W.); Chongqing Talent Plan Innovation and Entrepreneurship Team (2022, to E.L.); the New Cornerstone Science Foundation (to X.W.); the start-up grants of Wenzhou Institute, University of Chinese Academy of Sciences (No. WIUCASQD2025069 to J.M.); and Zhejiang Key Laboratory of Soft Matter Biomedical Materials (2025ZY01036, 2025E10072). We thank the staff at the Center for Biological Imaging (CBI), Institute of Biophysics (IBP), Chinese Academy of Sciences (CAS) for their technical support on the cryo-ET/EM, and we are grateful to Dr. Xiaojun Huang, Dr. Xujing Li and Dr. Boling Zhu for help with cryo-ET/EM data screening and collection. We also thank the high-performance computation platform of CBI, IBP, CAS for the computing resources.

## AUTHOR CONTRIBUTIONS

X.W., E.L., and L.X. supervised the project; J.M. prepared, screened, collected, and processed the cryo-ET sample and data; L.X. solved the structure of dimer of trimers of preF from RSV by sub-tomogram averaging, J.M. solved other RSV and hMPV cryo-ET structures by sub-tomogram averaging; J.M., X.W. and H.Z. analyzed the structures; H.Z. performed the functional assay, and analyzed the data; W.Y. expressed, purified the protein samples and solved the cryo-EM structures, refined and validated the atomic models; X.W., J.M., H.Z., W.Y., L.X. and E.L. wrote the manuscript with contributions from all authors. J.W., J.D., L.W., and R.F. assisted during sample making, data collection and analysis.

## DECLARATION OF GENERATIVE AI AND AI-ASSISTED TECHNOLOGIES IN THE WRITING PROCESS

During the preparation of this work, the authors used ChatGPT 5.6 for enhance grammar and readability during the manuscript writing. The authors reviewed and edited the output as needed and take full responsibility for the content of the published article.

## EXPERIMENTAL MODEL AND SUBJECT DETAILS

### Cells and Viruses

HEp-2 cells (ATCC CCL-23) and Vero E6 cells were cultured in Dulbecco’s Modified Eagle Medium (DMEM) supplemented with 10% (v/v) fetal bovine serum (FBS). The RSV Long strain was kindly provided by National Institutes for Food and Drug Control. It was propagated in either HEp-2 cells in DMEM containing 2% (v/v) FBS and was finally purified by density gradient centrifugation. The hMPV A1 strain was kindly provided by Dr. Yao Zhao (Children’s Hospital of Chongqing Medical University, Chongqing, China). hMPV was propagated in Vero E6 cells with 0.00025% trypsin and 2% FBS.

## METHOD DETAILS

### Cryo-ET sample preparation

Viruses (RSV or hMPV) were harvested directly from infected HEp-2 or Vero E6 cells cultured on gold Quantifoil grids (300 mesh, R1.2/1.3; Quantifoil MicroTools GmbH, Germany). The procedure generally followed established protocols for correlated cryo-electron tomography of mammalian cells ^52^. Briefly, grids were glow-discharged and subjected to ultraviolet irradiation for 30 min in a biosafety cabinet. After placing grids in a 6-well plate, cells were seeded at ∼30% confluence and allowed to adhere for 2–8 h in a humidified incubator (37 °C, 5% CO_2_). Adherent cells were then infected at a multiplicity of infection (MOI) of 1 and further incubated until widespread cytopathic effect was evident. For certain experiments, infected cells were treated with 5 mM EGTA and EDTA for 0, 0.5, or 2 days. In some hMPV samples, Fab fragments targeting the pre-F protein were also incubated with the virions for 30 min prior to freezing. Immediately before plunge-freezing, 3 μL of PBS was applied to the front side of each grid, and 5 nm BSA-coated gold fiducials (Electron Microscopy Sciences) were pipetted onto the back side. Grids were blotted from the back for 4 s using Leica EM GP2 or manually, and then vitrified in liquid ethane. All samples were inactivated before imaging. Vitrified grids were stored in liquid nitrogen until screening or data collection.

### Cryo-ET data collection and tomogram reconstruction

Initial sample screening was performed on a 200 kV Talos 200C transmission electron microscope equipped with a CCD camera (4096 × 4096 pixels). Whole-grid montages were acquired at 510 × magnification under high underfocus (∼8 mm) to visualize viral particles. Grid squares exhibiting a high density of virions within vitreous ice of suitable thickness were selected for high-resolution cryo-ET. Data collection was carried out on a 300 kV Titan Krios G3 microscope (Thermo Fisher Scientific) equipped with a post-column energy filter (GIF, Gatan) and a K2 Summit direct electron detector (Gatan Inc.). Tilt-series were recorded in super-resolution mode at pixel sizes of 1.36 Å or 2.73 Å. The energy filter slit was set to 20 eV, aligned to the zero-loss peak, and remained active throughout acquisition. Each tilt image was recorded as a 10-frame movie with an exposure dose of 3.5 e⁻/Å^2^ per tilt. Tilt-series were acquired automatically using SerialEM ^53^ over an angular range of –60° to +60° with a 3° increment, following a dose-symmetric scheme. Each series comprised 41 images, resulting in a total cumulative dose of approximately 143.5 e⁻/Å^2^. The defocus range was set between –2 and –5 μm.

All movie frames were aligned and summed using MotionCor2 ^54^ at bin 2. Tilt-series alignment and tomogram reconstruction were performed using IMOD v.4.10.38 (batchruntomo) ^55^ or AreTomo ^56^, with series exhibiting excessive stage drift or alignment errors >0.6 nm excluded. Final tomograms were reconstructed using the simultaneous iterative reconstruction technique (SIRT). Selected tomograms were further processed with IsoNet ^32^ at bin-10 (1.36 Å pixel) or bin-5 (2.73 Å pixel) to correct for missing-wedge artifacts and enhance signal-to-noise ratio for visualization. Three-dimensional visualization and analysis were performed using IMOD v.4.10.38 ^55^, and UCSF ChimeraX v.1.4 ^57^. A summary of reconstruction statistics is provided in Table S1.

### Sub-tomogram averaging of RNP complexes

Subtomogram averaging of viral RNP complexes was performed using emClarity v.1.6.1 ^58^. Initial particle picking was conducted on bin5 tomograms, using a single reference subtomogram derived from IsoNet-processed, missing-wedge-corrected patch ^32^. Candidate particles from each tomogram were visually inspected in IMOD v.4.10.38 ^55^ and validated using the Place Object plugin ^59^, during which process the false positives were manually excluded.

Particle coordinates were then imported into RELION v.4.0 ^60^ for iterative alignment and reconstruction. An initial round of alignment was performed under C1 symmetry to estimate helical parameters. Subsequent helical reconstruction employed the following constraints: inner diameter 5 nm, outer diameter 22 nm, twist –35 ± 2°, rise 7.5 ± 0.4 Å, with 6 unique asymmetric units and a central Z length of 25%. Following helical 3D refinement, misaligned particles were manually curated and removed using the Place Object plugin ^59^. Final reconstructions of RSV and hMPV RNPs achieved global resolutions of 10.2 Å and 10.9 Å, respectively. A summary of reconstruction statistics is provided in Table S2.

### Subtomogram averaging of RSV preF protein dimer of trimers

Raw tilt-series were processed in Warp v.1.0.9 for motion correction, defocus estimation, tilt-series sorting and dose-weighting ^61^. Visibly obstructed or out-of-field images were excluded. Tilt-series were aligned in AreTomo v.1.3.4 ^56^, and IMOD-compatible alignment files (.xf format) ^55^ were imported into Warp for tomogram reconstruction (231 tomograms in total). Template matching was performed at 8.16 Å (bin 6) in pyTOM v.1.1 ^62^ using a pre-F trimer reference (EMD-2392) ^21^. Initial reference was aligned and refined with emClarity v.1.6.1 ^58^. Particles located inside the viral envelope, on ice contaminants, near carbon edges, or duplicated were removed. A total of 171,304 subtomograms were extracted in Warp (box size 80 voxels, 3.6 Å/voxel). After iterative refinement and classification in RELION v3.0.8 and v4.0.1 ^60^, 40,554 particles containing a neighboring trimer were selected. Refinement under C1 symmetry yielded a 9.4 Å map. Following recentering at the dimer interface and re-extraction, refinement focusing on the region outside the membrane produced maps at 8.2 Å (C1) and 6.9 Å (C2). As no major differences were observed, the C2-symmetric reconstruction was used for further analysis. Fourier-shell correlation (FSC) calculation, resolution estimation and B-factor sharpening were performed in RELION; Noise2Noise-based denoising was applied in M ^61^. Statistics are summarized in Table S2.

### Subtomogram averaging of RSV preF protein of hexagonal array

Tilt-series with well-aligned hexagonal arrays were processed in emClarity v.1.6.1 ^58^. Particle picking was conducted at bin 5 using a single IsoNet-corrected subtomogram patch as a reference (C1) ^32^. Particles were visually validated in IMOD v.4.10.38 ^55^ and Place Object ^59^, during which process the false positives were manually removed. Given the clear hexagonal lattice in Fourier space, C3 symmetry was imposed during alignment in emClarity. Particles were then transferred to RELION v4.0 for further reconstruction ^60^. Two reconstruction strategies were employed: (1), Multi-trimer in presence of envelope: Alignment of a larger volume containing multiple trimers and the lipid envelope yielded maps at 11.0 Å (untreated) and 15.9 Å (treated with EGTA & EDTA for 2 days). (2), Single trimer only: Alignment focused solely on the protein region produced higher-resolution maps at 7.4 Å (untreated) and 9.0 Å (treated with EGTA & EDTA for 2 days). Reconstruction statistics are provided in Table S2.

### Sub-tomogram averaging of hMPV preF protein

Tilt-series with satisfactory alignment were processed using emClarity v.1.6.1 ^58^. Due to the less ordered arrangement of surface proteins on hMPV compared to RSV, direct reference picking from subtomograms was not feasible. Instead, template matching was performed at bin 5 using the RSV pre-F trimer structure (EMD-2392) as an initial reference ^21^, followed by alignment refinement within emClarity. The strong envelope signal in the template facilitated the localization of envelope-associated particles. Candidate particles from each tomogram were visually inspected in IMOD v.4.10.38 ^55^ and the Place Object plugin ^59^, during which process those clearly not associated with the viral envelope were manually excluded. Particle alignment was iteratively refined at bin 3 and then bin 2 using a box size of 25 nm × 25 nm × 20 nm, followed by multiple rounds of classification focused on the central protein density to further eliminate false positives. This process yielded two distinct classes corresponding to pre-F monomers and trimers.

Particles belonging to each class were pooled and transferred to RELION v4.0 ^60^ for final reconstruction. In the absence of Fabs, the pre-F monomer and trimer were resolved at 32.4 Å and 32.1 Å, respectively. In complexes with CNM3035 or CNM3005 Fabs, both monomeric and trimeric populations were observed. Monomeric pre-F was reconstructed in complex with a single Fab, whereas trimers were resolved with two bound Fabs. The final resolutions for Fab-bound complexes were as follows: CNM3035 Fab: monomer 29.2 Å, trimer 31.1 Å; CNM3005 Fab: monomer 26.6 Å, trimer 27.7 Å. A summary of reconstruction statistics is provided in Table S2.

### Functional assay

RSV Long strain and hMPV A1 strain were propagated in HEp-2 and Vero E6 cells. When cytopathic effect (CPE) reached approximately 90%, cell culture supernatants containing viral particles were harvested. For the EDTAand EGTAtreated assays, each virus preparation was divided into two groups: experimental group (E) and control group (C). The E group was treated with EDTA (5 mM) and EGTA (5 mM) for various durations (e.g., 0.5, 2, 3 days), while the C group remained untreated. At the designated time points, the E group was neutralized by addition of Mg^2+^ (5 mM) and Ca^2+^ (5 mM), whereas the C group was supplemented with a pre-neutralized mixture (containing 5 mM EDTA, 5 mM EGTA, 5 mM Mg^2+^, and 5 mM Ca^2+^). Subsequently, both groups were diluted 5-fold with 2% FBS-containing medium.

Then, Virus samples were aliquoted into 1.5 mL microcentrifuge tubes and heated in a metal block heater at various temperatures (37, 45, 50, 53, 55, 57, and 59 °C) for 10 minutes. Immediately after heating, samples were placed on ice for rapid cooling. Viral titers were subsequently determined using a modified plaque assay as followed.

One day before infection, HEp-2/Vero E6 cells were plated into 96-well plates at 2×10^4^ cells per well and maintained overnight at 37 °C in a humidified incubator with 5% CO_2_. On the day of infection, the growth medium was aspirated and replaced with 80 µL of the test virus sample per well to initiate viral infection. Each virus sample was tested in duplicate (two technical replicates) to ensure reproducibility. Then the plates were incubated at 37 ℃ for 1 hour. The cells were next overlaid with 150 μL of DMEM medium supplemented with 1.2% methylcellulose and 2% FBS. After incubation at 37 ℃ with 5% CO_2_ for 40 hours, add 100 μL of fixation buffer (4% paraformaldehyde) to fix cells at room temperature for 30 min. For hMPV A1, plaque-forming unit equivalents (PFU-Eq/mL) were directly visualized and quantified using a high-content imaging system (Opera Phenix) based on GFP fluorescence. In addition, PFU-Eq/mL were independently determined by enzyme-linked immunospot (ELISPOT) assay and showed good concordance with the GFP-based quantification. For RSV Long and hMPV A1, the modified plaque assay was conducted according to the protocol described below: Prepare 2% bovine serum albumin (BSA) for blocking and incubate at room temperature for 30 min. Then, add 50 μL per well of the RSV F-specific antibody solution incubating at room temperature for 1 hour and wash plate 3 times with filtered PBST (0.05% Tween-20). Next, add 50 μL per well of the secondary antibody (HRP goat anti-human IgG) solution and incubate at room temperature for 1 hour. Wash plate 3 times with filtered PBST (0.05% Tween-20). Chromogenic reactions were performed using TrueBlue (KPL) for visualization of RSV-infected plaques. Plates were scanned and biospots were counted by ImmunoSpot Analyzer (CTLAnalyzerLLC, USA) for virus titer determination.

### Expression and purification of hMPV pre-F protein and monoclonal Fabs

The pre-F protein of hMPV strain A1 (residues 1-490) containing the stabilizing mutations A185P, A113C, and A339C was used in this study ^13,63^. The construct included a C-terminal T4 fibritin trimerization motif followed by an 8×His tag and was cloned into the pCAGGS vector for mammalian expression.

For protein production, the plasmid was transiently transfected into suspension-adapted HEK293F cells cultured at 37 °C with 8% CO_2_ under constant agitation (130 rpm). After 72 hours post-transfection, cells were harvested by centrifugation (1000× g, 30 min), and the supernatant was collected. The secreted protein was first purified by nickel-affinity chromatography (Ni-NTA resin) and subsequently subjected to size-exclusion chromatography with a Superdex 200 column (GE Healthcare) in buffer containing 20 mM Tris-HCl pH 8.0 and 200 mM NaCl.

The monoclonal Fab fragments used in cryo-EM analyses were generated essentially as previously described ^63^. Briefly, monoclonal antibodies were screened against the stabilized pre-F trimer described above, and corresponding Fabs were prepared by papain digestion followed by standard affinity purification.

### Cryo-EM sample preparation and data collection

Purified hMPV pre-F trimer (3.0 mg/mL) was incubated with a 1.2-fold molar excess of each Fab (CNM3035 or CNM3005) in buffer containing 20 mM Tris-HCl pH 8.0 and 200 mM NaCl at 4 °C for 30 min. Prior to vitrification, 0.05% (w/v) n-dodecyl-β-D-maltoside (DDM) was added to the complex. A 3 μL aliquot of each sample was applied to a glow-discharged holey-carbon gold grid (C-flat, 300 mesh, R 1.2/1.3). Grids were blotted for 5 s with zero blot force under 100% humidity at 4 °C and plunge-frozen in liquid ethane using an FEI Mark IV Vitrobot. Data were collected on either a 200 kV Arctica or a 300 kV Titan Krios transmission electron microscope equipped with a K2 or K3 direct electron detector operating in counting mode. Movie stacks were recorded using SerialEM ^53^ with a defocus range of-0.5 to-3.0 μm.

### Cryo-EM data processing and 3D reconstruction

Motion correction of movie stacks and contrast transfer function estimation were performed in cryoSPARC v4.7.1 ^64^ using patch-based algorithms. Initial particle picking was carried out by a combination of blob picking and template matching with a reference pre-F trimer. Multiple rounds of 2D classification were applied to remove non-particle and contaminant images. Subsequent 3D processing involved iterative cycles of ab initio reconstruction, heterogeneous refinement, homogeneous refinement, and non-uniform refinement, yielding distinct classes corresponding to pre-F trimers and monomers in complex with Fab. For certain complexes, local refinement that excluded the C-terminal regions of the Fab was employed to further improve the local resolution of the interface between F-protein and the Fab. Final reconstruction statistics are summarized in Table S3.

### Atomic modeling, refinement, and validation

The atomic model of hMPV pre-F was derived from the previously reported structure (PDB 7SEJ). Fab N-terminal variable domains were modeled using AlphaFold3 ^65^. Initial rigid-body docking of the pre-F and Fab models into the cryo-EM density map was performed in UCSF ChimeraX v1.4 ^57^. Subsequently, the pre-F and Fab components were combined into a single composite model and manually adjusted in Coot v0.981 ^66^ to optimize fit and connectivity. Iterative rounds of real-space refinement in Phenix v1.14 ^67^ and manual rebuilding in Coot were carried out to improve model geometry and map correlation. Final models were validated using MolProbity ^68^ to assess stereochemical quality. All structural figures were prepared with UCSF ChimeraX v1.4 ^57^.

