## Supplementary material for "Architectural trade-offs between environmental stability and genomic redundancy reveal divergent pneumoviral entry strategies": SI figures 1-6 and tables 1-3

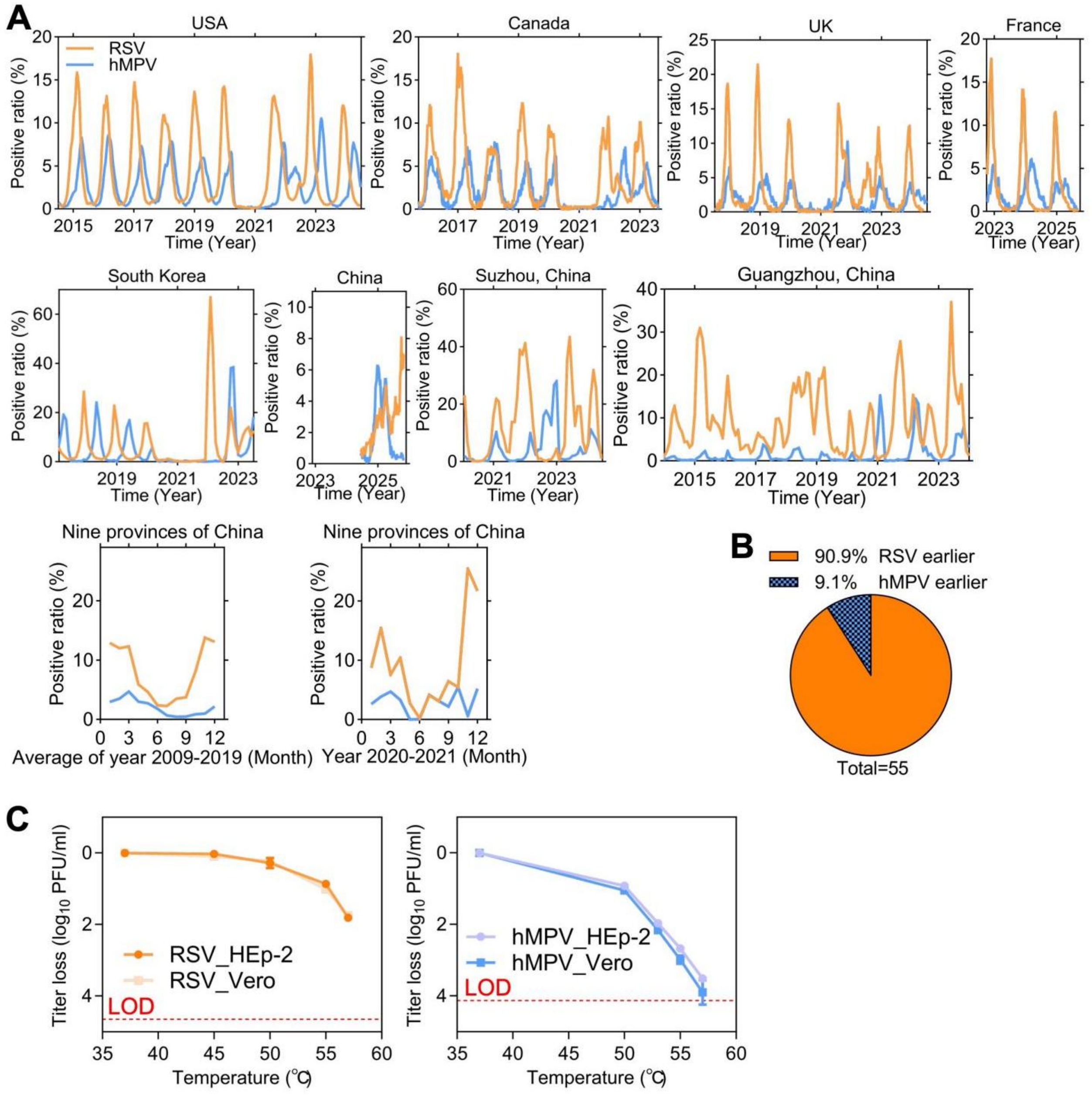


**Figure S1. Differential virion thermotolerance is associated with the seasonal separation of RSV and hMPV, related to Figure 1.**

(A) Multi-year surveillance time series showing RSV and hMPV detections across countries and regions. The surveillance data were compiled from publicly available respiratory-virus surveillance databases and published epidemiological studies.

(B) Quantification of the relative seasonal onset of RSV and hMPV outbreaks across surveillance seasons. To compare epidemic timing along the seasonal temperature transition, each surveillance season was defined from summer through the following spring. Outbreak onset was defined as the first time point at which detections reached 50% of the seasonal maximum for each virus. The pie chart shows the fraction of country/region-year observations in which RSV or hMPV emerged earlier.

(C) Infectious-titer loss of RSV and hMPV after 10 min of incubation at the indicated temperatures. Incubation at 37°C was used as the low-temperature reference condition. Data are mean ± SD.

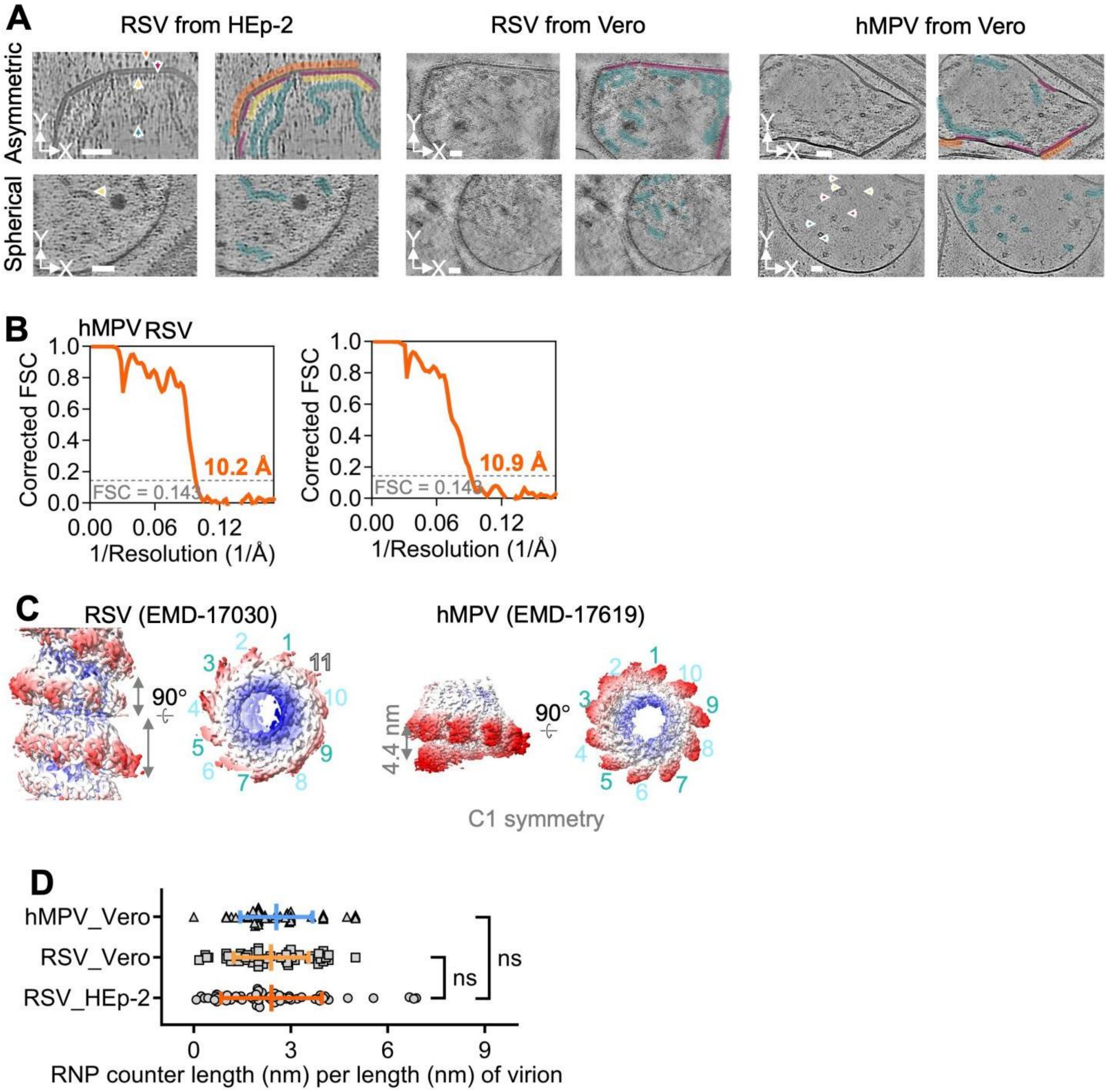


**Figure S2. Tomogram slices and statistics of the viruses, related to Figure 1.**

(A) Typical cryo-electron tomographic slices of released viruses. The G/F proteins outside the envelop is highlighted in orange, the M protein inside the envelop is highlighted in purple, the less well-ordered protein nearby the M protein was previously hypothesised being M2-1 is highlighted in yellow, and the helical RNP is highlighted in blue. Scale bars, 50 nm.

(B) The fourier shell correlation (FSC) plot of RNP structures from RSV and hMPV reconstructed by sub-tomogram averaging.

(C) Structure of in vitro RNP of RSV and hMPV. Both the full length RSV and hMPV are not canonical helical structure.

(D) The statistics of the RNP length per length of virion in filamentous virus particles. The error bars are standard divisions. Statistical significance was determined by two tailed unpaired t-test analysis.

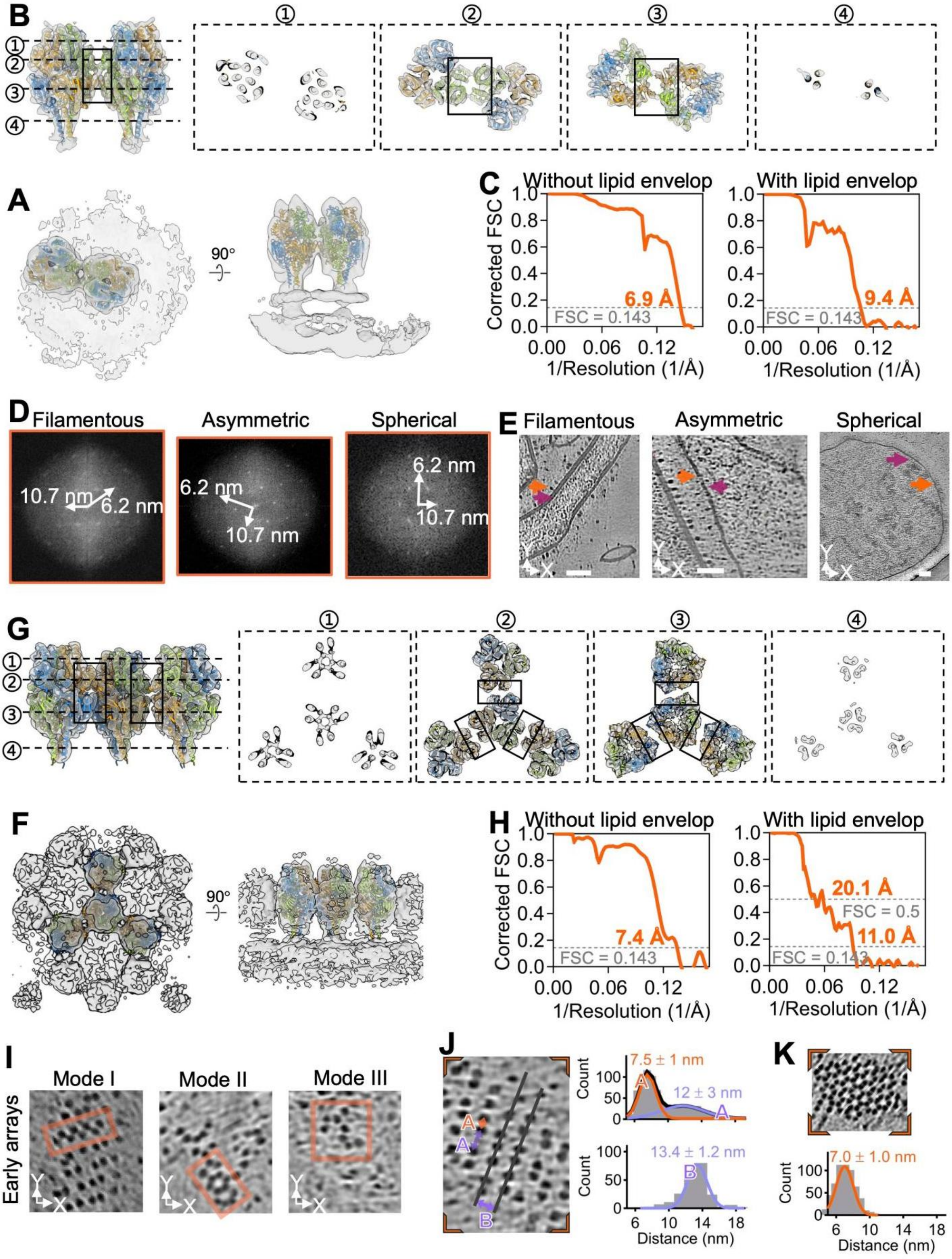


**Figure S3. Analysis of the cryo-electron tomographic maps and slices of RSV F protein, related to Figure 2.**

(A and B) Model fitting details (in orange, green, and blue) of the sub-tomogram averaging (in light gray) of the dimer of F trimers. The dimer interface details are highlighted in black boxes. The signal of lipid envelop was

excluded for better reconstruction in B, while was excluded to visulize the lipid envelop in A.
(C) The fourier shell correlation (FSC) plot of the maps of dimer F proteins.
(D) Fourier transforms of the array regions of the G/F proteins in Fig. 2B, indicating a hexagonal lattice with a 10.7 nm spacing (white arrow).
(E) Central slices of the cryo-ET of RSV particles. The positions of G/F protein and M protein are highlighted with orange and purple arrows, although the proteins maybe absence in the micrograph. Scale bars, 50 nm.
(F and G) Model fitting details (in orange, green, and blue) of the sub-tomogram averaging (in light gray) of the hexagonal array of F trimers. The dimer interface details are highlighted in black boxes. The signal of lipid envelop was excluded for better reconstruction in E, while was excluded to visulize the lipid envelop in D.
(H) The fourier shell correlation (FSC) plot of the maps of hexagonal array of F protein.
(I) Examples of early arrays of F protein. The error bars are standard divisions (SD).
(J) Zoom in of cryo-electron tomographic surface slice of the major filamentous RSV in Fig. 2A. The distance between two nearby G/F proteins in the same column was denoted as A and showed two gaussian peaks, while the distance between two nearby G/F proteins in the nearby column was denoted as B and showed only one gaussian peak. The error bars are standard divisions (SD).
(K) Zoom in of cryo-electron tomographic surface slice of the minor filamentous RSV in Fig. 3B. The distance between two nearby G/F proteins showed only one gaussian peak.

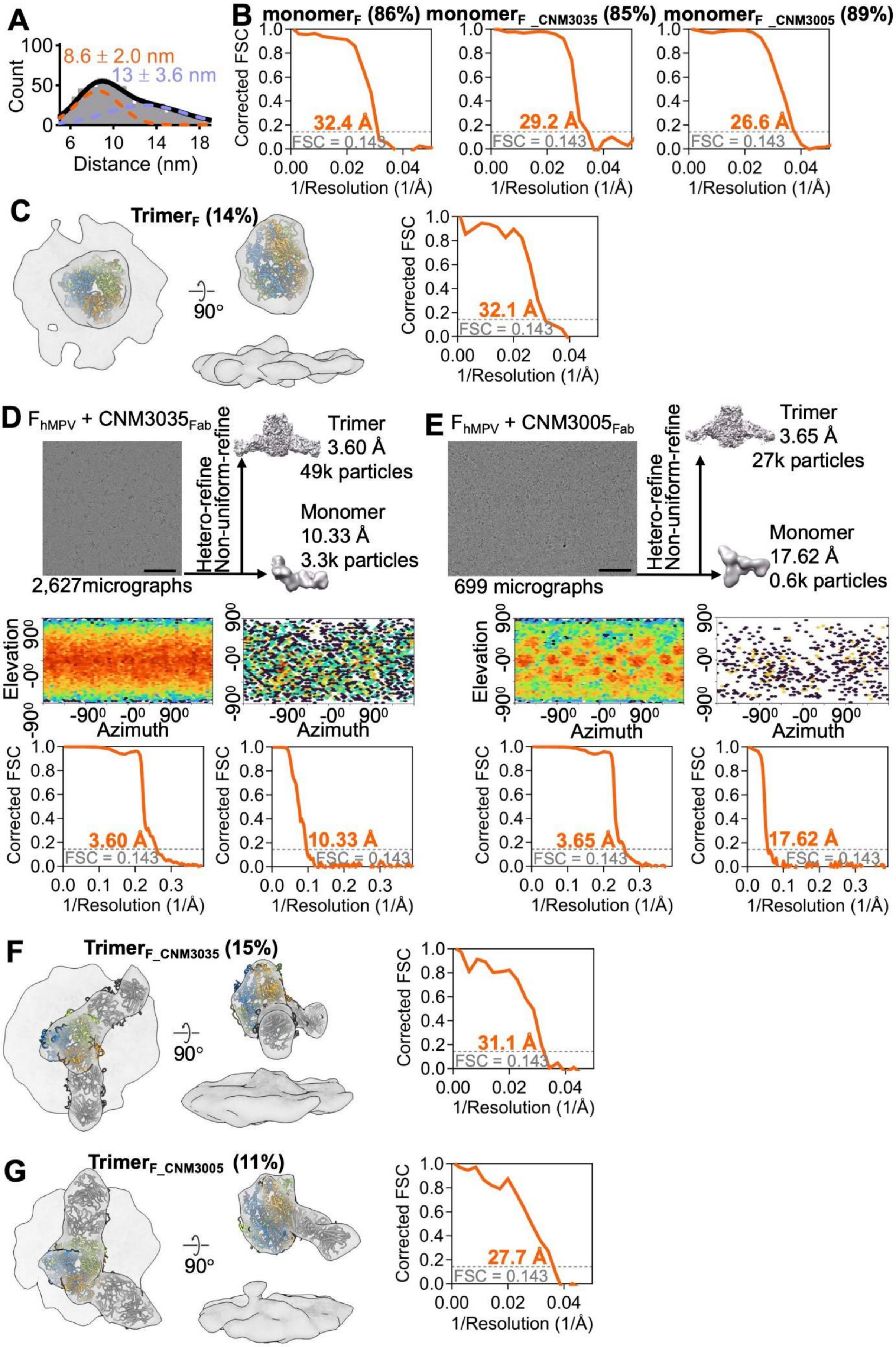

A
Count
8.6 ± 2.0 nm
13 ± 3.6 nm
Distance (nm)
B
monomerF (86%)
monomerF _CNM3035 (85%)
monomerF _CNM3005 (89%)
Corrected FSC
32.4 Å
29.2 Å
26.6 Å
FSC = 0.143
1/Resolution (1/Å)
C
TrimerF (14%)
90°
32.1 Å
D
FhMPV + CNM3035Fab
2,627micrographs
Hetero-refine
Non-uniform-refine
Trimer
3.60 Å
49k particles
Monomer
10.33 Å
3.3k particles
Elevation
Azimuth
3.60 Å
10.33 Å
E
FhMPV + CNM3005Fab
699 micrographs
Trimer
3.65 Å
27k particles
Monomer
17.62 Å
0.6k particles
3.65 Å
17.62 Å
F
TrimerF_CNM3035 (15%)
31.1 Å
G
TrimerF_CNM3005 (11%)
27.7 Å

**Figure S4 Analysis of the cryo-electron tomographic maps of hMPV F protein, related to Figure 3.**

(A) The distance between two nearby G/F proteins of hMPV particles. The distances could be fitted with two gaussian peaks. The error bars are standard divisions (SD).

(B) The FSC plots of the sub-tomogram averaging of hMPV F monomers in absence of Fab or in presence of CNM3035 or CNM3005 Fab fragments.

(C) Surface representations of the sub-tomogram averaged maps overlapped with the corresponding structure models of hMPV pre-F trimer (map in light gray, model in orange, green and blue). The right panel shows the corresponding FSC plots of the sub-tomogram averaging map.

(D and E) Cryo-EM data processing of the hMPV F protein in complex with CNM3035 and CNM3005 Fab fragments. Scale bars, 100 nm.

(F and G) Surface representations of the sub-tomogram averaged maps overlapped with the corresponding structure models of hMPV pre-F trimer (map in light gray, model in orange, green and blue) in complex with two CNM3035 (d) or two CNM3005 Fab fragment (map in light gray, model in dark gray). The right panel shows the corresponding FSC plots of the sub-tomogram averaging maps.

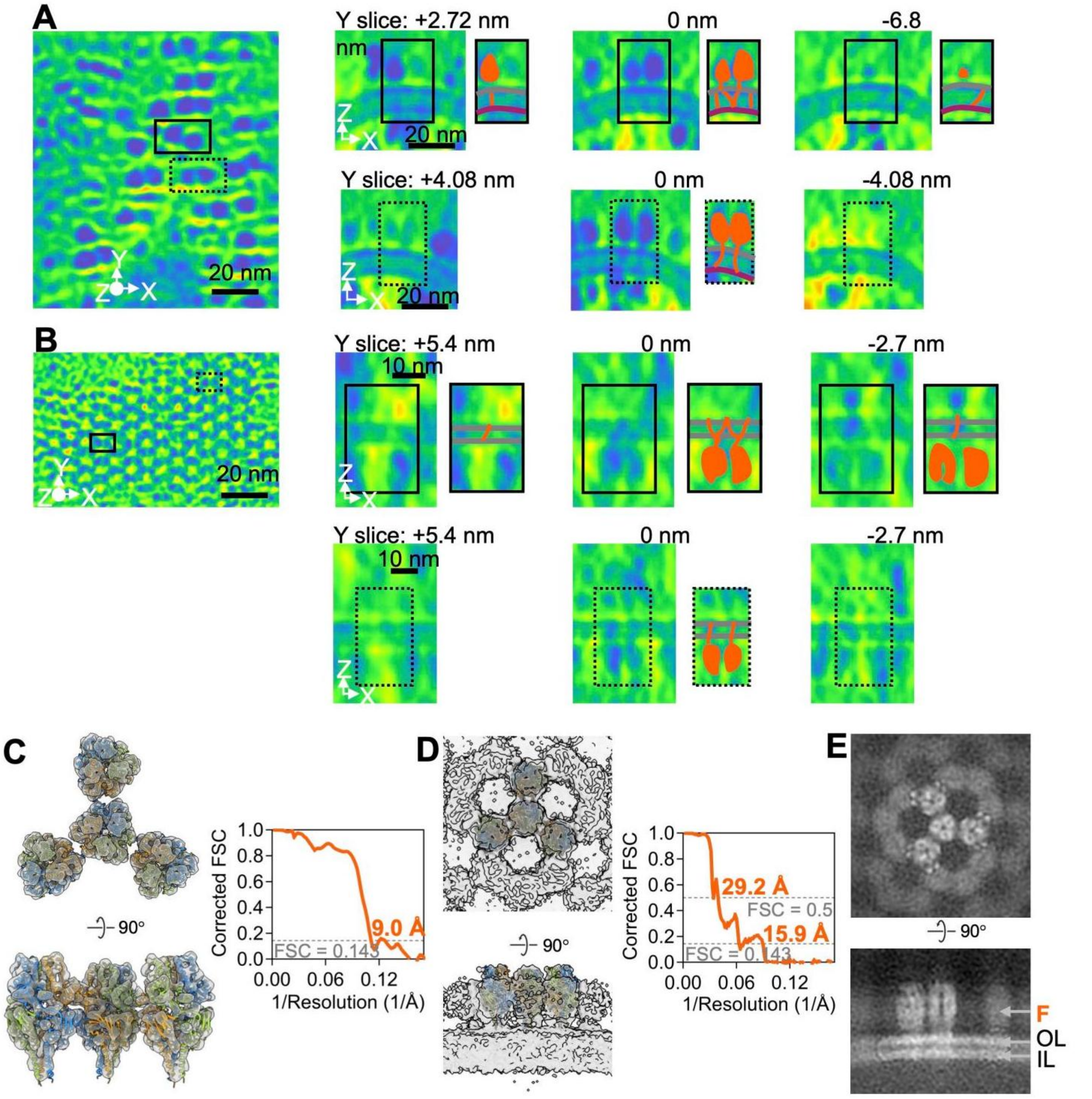


**Figure S5. Analysis of the cryo-electron tomography of the transmenbrance segment of RSV F in dimers and lattice, related to Figure 4.**

(A and B) Typical cryo-electron tomographic surface slices of RSV showing the TM-CT regions of pre-F in dimers and lattice. Both typical non-trimeric and trimerical are shown. For better ilustration, the F proteins are in orange, the M protein inside the envelop are in purple, the viral membrane is in gray. The thickness of the slices is 2.72 nm.

(C-E) Analysis of the cryo-electron tomographic maps of RSV F protein after treatment of EDTA/EGTA for 2 days. Model fitting (in orange, green, and blue) of each monomer in the sub-tomogram averaging (in light gray) with the F trimers (C). The signal of lipid envelop was excluded for better reconstruction. Model fitting (in orange, green, and blue) of the sub-tomogram averaging (in light gray) of the hexagonal array of F protein (D). The signal of lipid envelop was included to visulize the lipid envelop. (E) Typical slices of the sub-tomogram averaging of the hexagonal arrays of F protein reconstructed as in E. The signal of M protein is abssent inside the outer leaflet (OL) and inner leaflet (IL) of lipid bilayer.

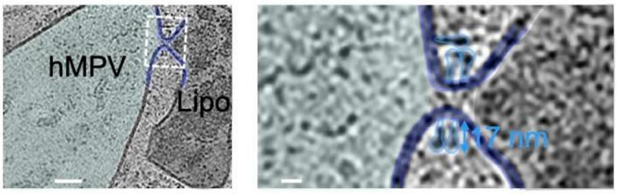


**Figure S6. Fusion-associated intermediates captured on a hMPV particle, related to Figure 5.**
Representative tomographic slice showing an hMPV particle with a fusion-pore-like membrane-continuity event. The boxed region is enlarged on the right to highlight associated F-like densities.

**Table S1. Summary of cryo-ET data collection.**

| Samples | RSV | RSV | RSV | RSV | hMPV | hMPV_CNM3035Fab | hMPV_CNM3005Fab | hMPV | hMPV |
|---|---|---|---|---|---|---|---|---|---|
| | HEp-2 | Vero | HEp-2 | | Vero | | | | |
| | 0.0 Day | 0.0 Day | 0.5 Day | 2.0 Day | 0.0 Day | 0.0 Day | 0.0 Day | 0.5 Day | 2.0 Day |
| **Voltage (kV)** | 300 | | | | | | | | |
| **Detector** | K2 | | | | | | | | |
| **Energy filter** | Bioquantum 20 eV slit | | | | | | | | |
| **Super-resolution mode** | Yes | | | | | | | | |
| **Defocus (μm)** | -2 to -5 | | | | | | | | |
| **Pixel size (Å)** | 1.36 & 2.73 | 2.73 | 1.36 | 2.73 | | | | | |
| **Acquisition scheme** | Dose-Symmetric, -60/60, 3º step, group 2 | | | | | | | | |
| **Total dose (e/Å2)** | 143.5 | | | | | | | | |
| **Frames per tilt** | 10 | | | | | | | | |
| **Number of tilt series** | 231 + 258 | 27 | 24 | 76 | 122 | 92 | 83 | 30 | 36 |

**Table S2. Summary of sub-tomogram averaging.**

| Samples | RSV + HEp-2 + 0.0 Day | | | RSV + HEp-2 + 2.0 Day |
|---|---|---|---|---|
| | RNP | $F_{Dimer\text{-}of\text{-}trimer}$ | $F_{Hexagonal\text{-}lattice}$ | $F_{Hexagonal\text{-}lattice}$ |
| **Final particles** | 24,527 | 40,554 | 15,280 | 5,628 |
| **Symmetry imposed** | Helical | C2 & C1 | C3 | C3 |
| **Map resolution (Å)** | 10.2 Å | 6.9 Å (C2, no lipid)<br>9.4 Å (C1, with lipid) | 7.4 Å (no lipid)<br>11.0 Å (with lipid) | 9.0 Å (no lipid)<br>15.9 Å (with lipid) |
| **FSC threshold** | 0.143 | | | |
| **Accession codes (EMD)** | 82792 | 82797<br>82796 | 82803<br>82800 | 82804<br>82806 |

| Samples | hMPV + Vero + 0.0 Day | | | hMPV_CNM3035$_{Fab}$ + Vero + 0.0 Day | | hMPV_CNM3005$_{Fab}$ + Vero + 0.0 Day | |
|---|---|---|---|---|---|---|---|
| | RNP | $F_{Monomer}$ | $F_{Trimer}$ | $F_{Monomer\text{-}CNM3035}$ | $F_{Trimer\text{-}CNM3035}$ | $F_{Monomer\text{-}CNM3005}$ | $F_{Trimer\text{-}CNM3005}$ |
| **Final particles** | 18,425 | 1,780 | 283 | 1,605 | 288 | 2,470 | 319 |
| **Symmetry imposed** | Helical | C1 | | | | | |
| **Map resolution (Å)** | 10.9 Å | 32.4 Å | 32.1 Å | 29.2 Å | 31.1 Å | 26.6 Å | 27.7 Å |
| **FSC threshold** | 0.143 | | | | | | |
| **Accession codes (EMD)** | 82793 | 82808 | 82807 | 82810 | 82809 | 82813 | 82811 |

**Table S3. Summary of cryo-EM data collection, refinement and validation statistics.**

| Samples | FhMPV + CNM3035$_{Fab}$ | FhMPV + CNM3005$_{Fab}$ |
|---|---|---|
| **Data collection and processing** | | |
| Voltage (kV) | 200 | 300 |
| Detector | Gatan K2 | Gatan K3 |
| Energy filter | Bioquantum 20 eV slit | |
| Electron exposure (e/Å$^2$) | 60 | |
| Defocus range (μm) | -0.5 to -2.0 | -0.5 to -2.5 |
| Pixel size (Å) | 1.0 | 1.07 |
| Number of frames | 32 | |
| Number of images | 2,627 | 699 |
| Symmetry imposed | C3 (trimer) | |
| Final particle images | 49k | 27k |
| Map resolution (Å) | 3.60 | 3.65 |
| FSC threshold | 0.143 | |
| **Refinement** | | |
| Initial model used | PDB: 7SEJ for F protein; Predicted model by Alphafold3 for antibody | |
| Model resolution (Å) | 3.83 | 3.74 |
| FSC threshold | 0.5 | 0.5 |
| Map sharpening *B* factor (Å$^2$) | 122.1 | 128.9 |
| Model composition | | |
| Non-hydrogen atoms | 14,436 | 14,677 |
| Protein residues | 1,939 | 1,958 |
| *B* factors (Å$^2$) | | |
| Protein | 10.24 | 31.15 |
| R.m.s. deviations | | |
| Bond lengths (Å) | 0.004 | 0.003 |
| Bond angles (°) | 0.739 | 0.722 |
| Validation | | |
| MolProbity score | 1.77 | 1.64 |
| Clashscore | 5.87 | 4.44 |
| Poor rotamers (%) | 0.89 | 1.00 |
| Ramachandran plot | | |
| Favored (%) | 93.03 | 93.62 |
| Allowed (%) | 6.97 | 6.38 |
| Disallowed (%) | 0.00 | 0.00 |
| Accession codes | EMD-82866<br>PDB 44RU | EMD-82865<br>PDB 44RT |
